# Breadth of Fc-mediated effector function delineates grades of clinical immunity following human malaria challenge

**DOI:** 10.1101/2022.10.11.511755

**Authors:** Irene N. Nkumama, Dennis Odera, Fauzia Musasia, Kennedy Mwai, Lydia Nyamako, Linda Murungi, James Tuju, Kristin Fürle, Micha Rosenkranz, Rinter Kimathi, Patricia Njuguna, Mainga Hamaluba, Melissa C. Kapulu, Roland Frank, CHMI-SIKA study Team, Faith H. A. Osier

## Abstract

Controlled human malaria challenge studies in semi-immune volunteers provide an unparalleled opportunity to robustly identify mechanistic correlates of protection. We leveraged this platform to undertake a head-to-head comparison of seven functional antibody assays that are relevant to immunity against the erythrocytic merozoite stage of *Plasmodium falciparum*. Fc-mediated effector functions were strongly associated with protection from clinical symptoms of malaria and exponential parasite multiplication while the gold standard growth inhibition assay was not. The breadth of Fc-mediated effector function clearly discriminated grades of clinical immunity following challenge. These findings present a paradigm shift in the understanding of the mechanisms that underpin immunity to malaria and have important implications for vaccine development.

## INTRODUCTION

The Fab-dependent neutralization of bacteria, viruses and parasites has long been considered the main mechanism of antibody function that leads to protective immunity (Forthal, 2014; Lu *et al*., 2018). Neutralization typically occurs when antibodies prevent the binding of pathogens to host cell receptors and thereby inhibit invasion. In contrast, antibody Fc-dependent effector functions are less well-studied but appear to be increasingly important in mediating protection against infectious diseases such as HIV (Su *et al*., 2019), Ebola (Gunn *et al*., 2018) and COVID-19 (Winkler *et al*., 2021).

Fab-dependent neutralization is the main functional readout of immunity against *Plasmodium falciparum* malaria. It is typically assessed via the growth inhibition assay (GIA) and is used extensively to evaluate, prioritize and quantify the efficacy of blood-stage vaccine candidates. This continues to be case even though it does not reliably predict either naturally acquired or vaccine-induced protection (Duncan, Hill and Ellis, 2012). Malaria vaccines inducing high levels of GIA have had limited success in clinical trials in humans (Duncan, Hill and Ellis, 2012; Douglas *et al*., 2015, 2019; Payne *et al*., 2016; Minassian *et al*., 2021).

A growing body of evidence suggests that Fc-mediated effector functions are also important for protection against malaria (Teo *et al*., 2016; Moormann, Nixon and Forconi, 2019). Antibodies inducing Fc-effector function against merozoite stage parasites are acquired with age and correlated with protection in independent cohort studies (Joos *et al*., 2010; Hill *et al*.,2013; Osier *et al*., 2014a; Boyle *et al*., 2015; Tiendrebeogo *et al*., 2015). However, each study examined a single cellular or soluble effector such as neutrophils (Joos *et al*., 2010; Feng *et al*., 2021), monocytes (Hill *et al*., 2013; Osier *et al*., 2014a), natural killer cells (Odera *et al*.,2021) or complement (Boyle *et al*., 2015; Reiling *et al*., 2019) in a distinct study population. Neither the breadth, nor the relative importance of individual Fc-effector functions have been analyzed concurrently in a well-characterized experimental malaria study in humans.

We therefore utilized a unique controlled human malaria infection (CHMI) study conducted in semi-immune Kenyan adults to address these gaps. This platform has several distinct advantages over standard cohort studies. First, the precise timing, strain and dose of the inoculum is known, and hence minimizes the misclassification bias that inevitably occurs when these parameters cannot be accounted for in prospective cohort studies (Kinyanjui *et al*.,2009). Second, the close monitoring of study volunteers in residential facilities enabled the accurate capture of important clinical outcomes as soon as they occurred following challenge. This early identification was vital for safety (Kapulu, Njuguna and Hamaluba, 2018; Kapulu *et al*., 2021), and helped to minimize the misclassification bias that arises due to incidental illnesses that are often presumed to be malaria in the community. Third, analyses dependent on the time-to-event such as Cox regression, could be conducted with a greater confidence since the timing of exposure was known, and the rapidity with which an endpoint was met directly reflected *in vivo* parasite growth. Consequently, the precision around the estimates of the correlates of protection is significantly higher than that observed in cohort studies (Kapulu *et al*., 2022).

We used this framework to test the largest panel of Fc-mediated functional assays targeting *P. falciparum* merozoites in a single study; antibody-dependent respiratory burst from neutrophils (Joos *et al*., 2010;), antibody-dependent complement fixation (AbC’, Boyle *et al*.,2015), antibody-mediated natural killer cells activation (Ab-NK, Odera *et al*., 2021), and opsonic phagocytosis of both merozoites (Osier *et al*., 2014a) and ring stage parasites (Musasia *et al*., 2022). We also measured the gold-standard growth inhibitory activity (Persson et al., 2006) and asked which of these seven assays best predicted the primary clinical outcome, defined as the need for anti-malarial treatment post challenge (Kapulu *et al*., 2021). In this context, protected individuals are those who did not require treatment while susceptible ones did. We further defined sub-groups within these two categories; protected volunteers were either negative for parasitemia by polymerase chain reaction throughout the study (PCR-) or were PCR+ but had densities below 500 parasites/μl. Susceptible volunteers developed a fever (temperature > 37.5°C at any point during the study) with accompanying parasitemia of any density or remained afebrile but parasitemia exceeded a predefined threshold of 500 parasites/μl. These subgroups are subsequently referred to as the clinical grades of immunity and range in order from the most to the least immune; a) PCR-, b) PCR+, c) afebrile with parasitemia > 500/μl and d) febrile with any level of parasitemia.

We found that each Fc-mediated effector function was a significantly stronger predictor of protection than the GIA. Study volunteers that had high levels of function across the six antibody Fc-mediated effector assays tested were completely protected and did not require treatment post challenge. These data represent a paradigm shift in the understanding of the mechanisms that underpin naturally acquired immunity to malaria and have important implications for the design and evaluation of highly effective vaccines.

## RESULTS

### Controlled human challenge provides clear endpoints for clinical immunity

To distinguish adults with protective naturally acquired immunity,142 Kenyan adults that had previously been exposed to malaria were intravenously challenged with 3,200 sporozoites and closely monitored for parasitemia and clinical symptoms for up to 21 days (**Figure 1A,** Kapulu *et al*., 2021). Protection was defined as a parasitemia of <500 parasites/μl coupled with a lack of fever or other clinical symptoms of malaria until day 21 post-challenge when the study was completed. Eighty six of one hundred and fourty two (86/142) volunteers were protected and either remained parasite negative by PCR (PCR-) or had patent parasitaemia (PCR+). They did not receive any treatment until day 21 (**Figures 1B** and **1C**). The remaining fifty six (56/142) either developed parasitemia >500/μl or became febrile, were treated immediately and classified as susceptible (**Figure 1D and 1E**). Thus, in addition to whether volunteers were protected or susceptible, four clinical endpoints formed the basis for the immunological comparisons; PCR-, PCR+, afebrile with parasitemia > 500/μl and febrile with parasitemia of any density.

**Figure 1:**
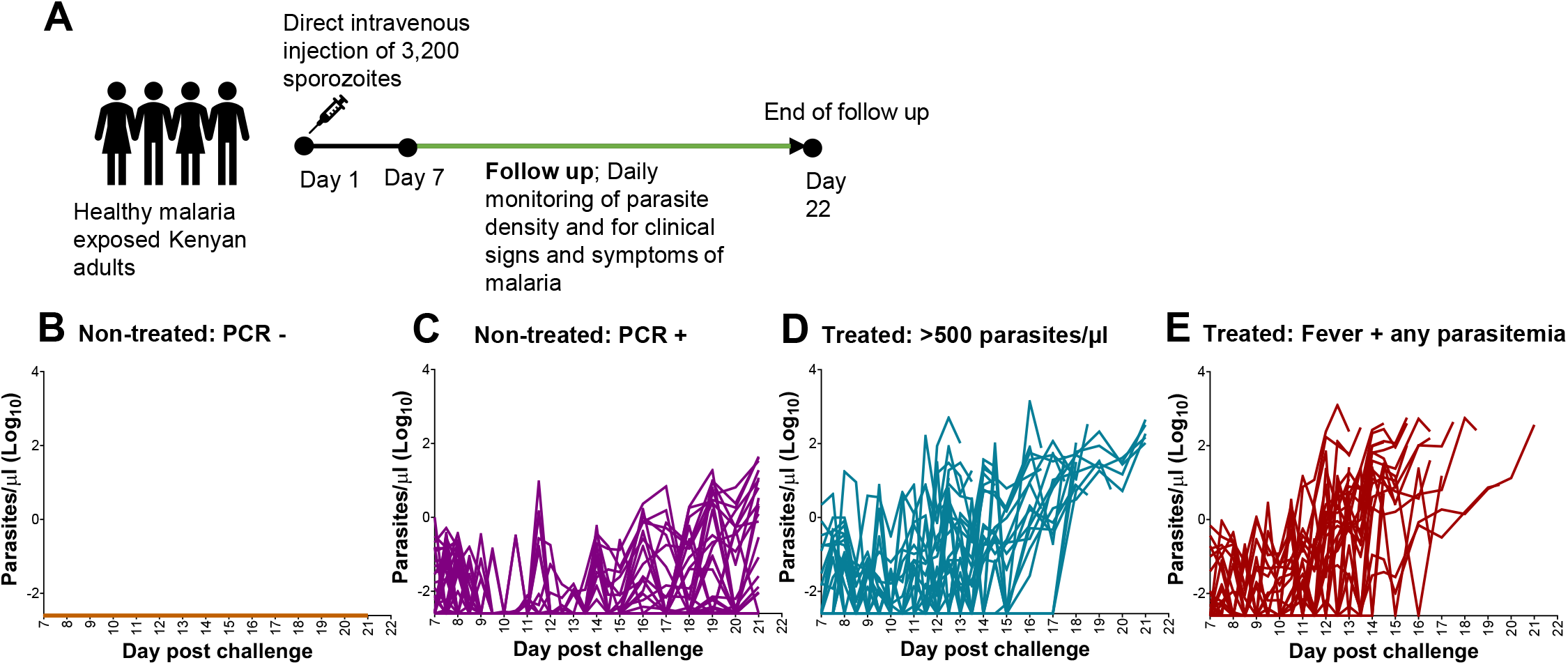
Controlled human challenge provides clear endpoints for clinical immunity. (**A**) Study design. Kenyan adult volunteers from geographical areas with varying levels of malaria transmission intensity were infected with 3200 live *P. falciparum* sporozoites via direct venous injection. Parasite densities were quantified by qPCR from day 7 to day 21. Volunteers were treated if parasitaemia exceeded >500/μl of blood or if they developed clinical signs and symptoms of malaria such as fever (axillary body temperature ≥37.5°C), chills, rigor, headaches, muscle weakness and fatigue. (**B-E**) Parasite densities during the follow up period for all 142 volunteers. A proportion were not treated until the end of the study on day 21 because they either (**B**) remained PCR-, n = 33/142 or (**C**) were PCR+ but parasitaemia was below the predefined threshold of 500/μl, n = 53/142. The remainder required treatment as (**D**) parasitaemia exceeded > 500/μl, n = 30 or (**E**) they became febrile, n = 26. Each line represents an individual. These primary data are published (Kapulu et al., 2020).

### Protection was associated with high levels of cytophilic antibodies against merozoites

To determine the role of antibodies against merozoites in NAI to malaria, we measured IgG, IgM and IgG subclasses in plasma samples collected a day before challenge. Surprisingly, total IgG and IgM antibodies against merozoites were equally abundant with a prevalence of 68% and 76%, respectively (**Figure 2A**). Consistent with previous studies, anti-merozoite IgG subclass antibodies were predominantly cytophilic IgG1 (74%) and IgG3 (59%), while non-cytophilic IgG2 (28%) and IgG4 (16%) antibodies were less common (**Figure 2A**). Protected volunteers were more likely to have anti-merozoite antibodies across all isotypes, than those who were susceptible (**Figure 2B**). This remained true within sub-groups, where PCR+ and PCR-volunteers had consistently higher anti-merozoite antibodies than those with parasitaemia >500/μl or fever (**Figure S1A**). In contrast, the prevalence of anti-tetanus toxoid IgG was similar across all sub-groups (**Figure 2B & Figure S1A**).

**Figure 2:**
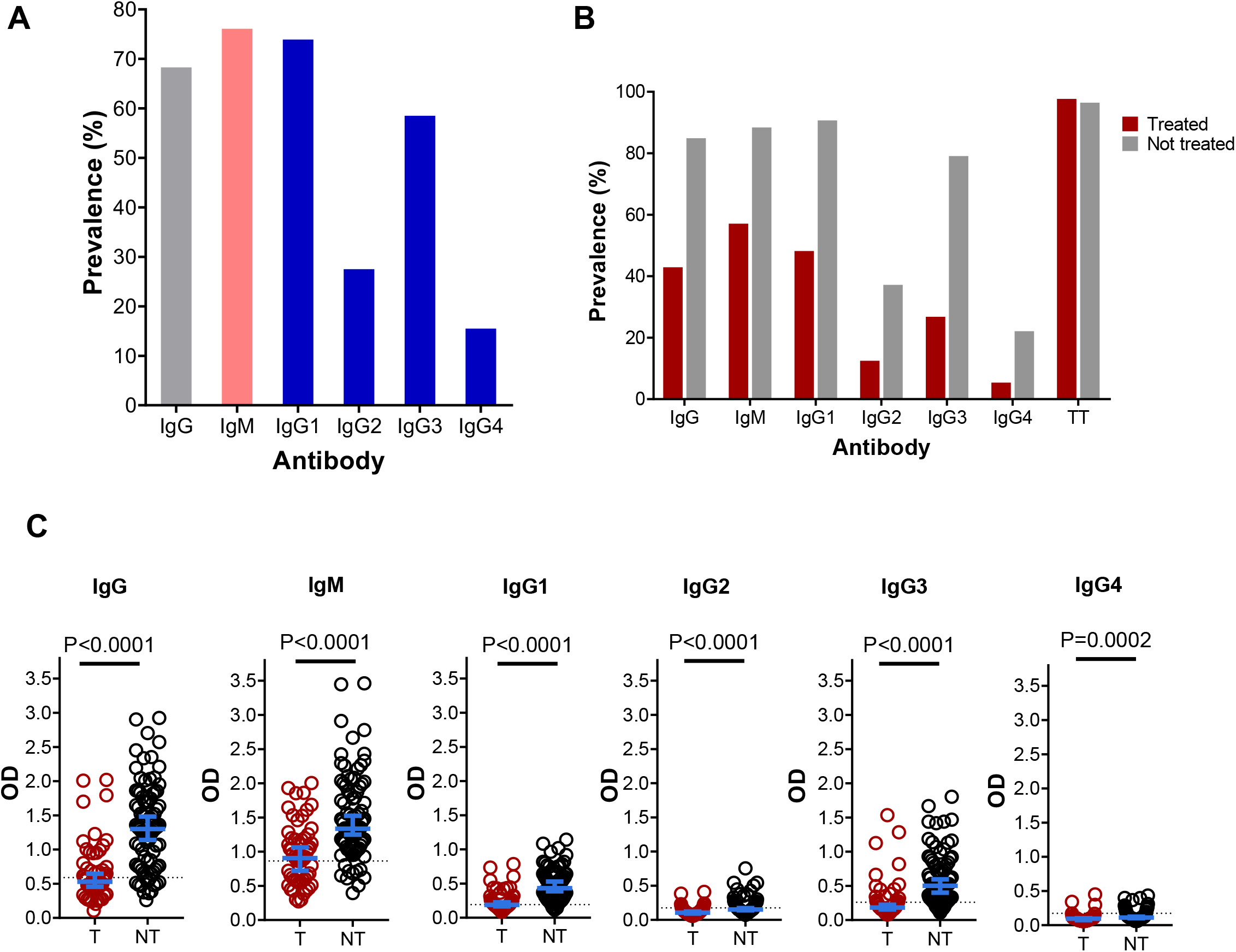
Protection was associated with high levels of antibodies against merozoites. (**A**) The overall prevalence of IgG, IgM, and IgG1-4 antibodies against merozoites. (**B**) The prevalence of IgG, IgM, and IgG1-4 against merozoites and total IgG against tetanus toxoid extract was compared between treated (n=56) and non-treated (n=86) volunteers. (**C**) IgG, IgM, and IgG1-4 antibodies levels compared between treated (T, n=56) and non-treated (NT, n=86) volunteers. Error bars represent median and 95% confidence intervals. p values were calculated using Mann-Whitney test. The dotted black horizontal line represents the seropositivity cut-off (mean + 3SD of malaria naïve plasma samples).

Anti-merozoite antibody levels followed the same pattern and were higher in protected versus susceptible volunteers (*p* < 0.001, **Figure 2C**). Similarly, PCR+ and PCR-volunteers had comparable antibody levels that were consistently higher than those detected in susceptible volunteers, and particularly those with fever (**Figure S1B**). Although both IgM and IgG discriminated the PCR+ and PCR-volunteers from those that were febrile, the effect was greater for IgG (**Figure S1B**). In the same vein, anti-merozoite IgG but not IgM discriminated febrile vs non-febrile volunteers (**Figure S1B**). Interestingly, despite the fact that non-cytophilic IgG2 and IgG4 antibodies were the least abundant, their levels nevertheless differed significantly within subgroups. However, these differences were less marked than those of the cytophilic antibodies. Finally, although levels of IgG to tetanus toxoid did not differ significantly at the subgroup level, in contrast to antibody prevalence, an interesting trend of increasingly higher levels in the most protected subgroups was observed (**Figure S1C**).

### Fc-mediated effector function was superior to neutralization in discriminating clinical immunity

The GIA assay is similar to neutralization and is the most widely used correlate of protection for blood stage malaria vaccine candidates (Duncan, Hill and Ellis, 2012). However, anti-merozoite antibodies have also been shown to recruit effector cells and complement (Moormann, Nixon and Forconi, 2019). We compared the ability of the GIA versus a panel of discrete Fc-mediated assays to distinguish between protected and susceptible volunteers in samples collected a day before challenge. We compared these immune measures further using the four endpoints of clinical immunity. For the GIA, we cultured blood stage parasites of the NF54 strain used in our challenge study for 96 hours (two cycles of parasite replication), in the presence of dialysed and heat-inactivated plasma (Persson et al., 2006). For Fc-effector function, we measured ADRB, AbC’, Ab-NK, and opsonic phagocytosis of merozoites and rings stage parasites. The Ab-NK assay had two readouts: degranulation (the proportion of NK cells that were CD107a^+^) and IFNγ production (the proportion of NK cells that were IFNγ^+^, Odera *et al*., 2021).

GIA levels were slightly higher in protected versus susceptible volunteers (*p* = 0.0156, **Figure 3A**). In marked contrast, each individual Fc-mediated effector function significantly distinguished protected versus susceptible volunteers (p < 0.001, **Figure 3A**). Similar results were obtained using the receiver operating characteristic (ROC) curves analysis (AUC=0.62 for GIA versus 0.69-0.80 for Fc-mediated functions, **Figure S2**). Additionally, the ability of the GIA to discriminate the four clinical endpoints was marginal (*p* = 0.0587, **Figure 3B**) compared to Fc-mediated effector functions which were consistently higher in PCR+ and PCR-volunteers compared to those that were febrile (**Figure 3B**).

**Figure 3:**
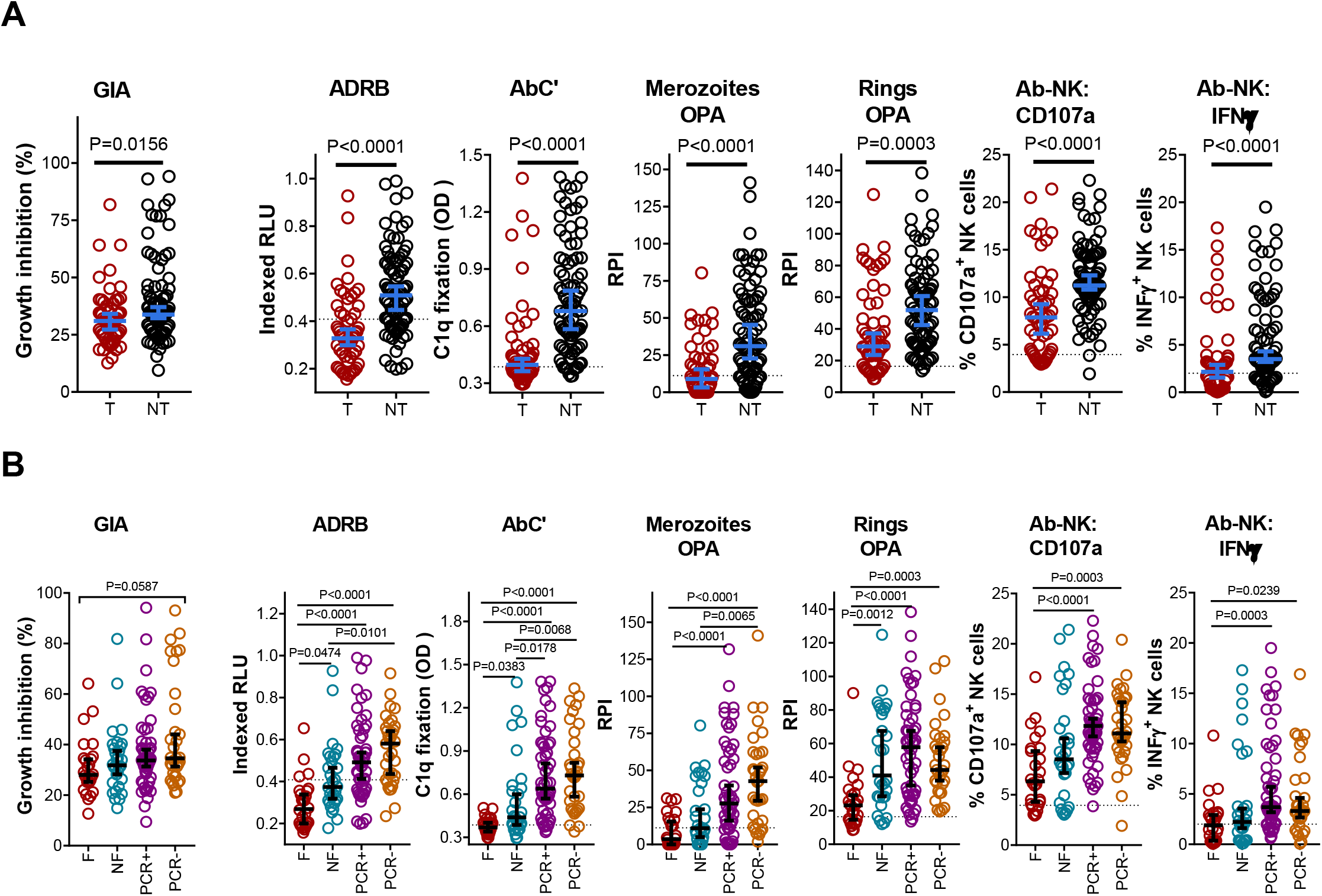
Naturally acquired anti-merozoite antibodies mediate protection via Fc-mediated effector function. **(A)** GIA (neutralization) and Fc-mediated effector functions were compared between all volunteers who were treated (T, n=56) and those who were not (NT, n=86). **(B)** GIA and Fc-mediated effector functions were compared across the four clinical endpoints; treated febrile (F, n=26), treated non-febrile (NF, n=30), not treated PCR+ (n=53) and not treated PCR- (n=33). Error bars represent the median and 95% confidence intervals. P values were calculated using Mann-Whitney test for treatment outcomes and using the Kruskal Wallis test with Dunn’s multiple comparisons test for the different phenotypes. The dotted black horizontal line represents the seropositivity cut-off (mean + 3SD of malaria naïve plasma samples). ADRB; antibody dependent respiratory burst by neutrophils, AbC’; complement fixation, OPA; opsonic phagocytosis by monocytes, ab-NK: Fc receptor-mediated natural killer (ab-NK) cell degranulation (CD107a) and IFNγ production, RLU; relative light units, RPI; relative phagocytosis index

### Fc-dependent function is correlated with anti-merozoite IgG and IgM but GIA is not

Next, we tested if the effector functions were correlated with the levels of anti-merozoite antibodies present in the volunteers prior to challenge. Each Fc-mediated effector function was more strongly correlated with total anti-merozoite IgG than IgM antibodies (**Figure 4A**). Not surprisingly, within the IgG subtypes, correlation coefficients for Fc-mediated effector functions were highest for cytophilic than non-cytophilic antibodies (r values 0.51 to 0.86 versus 0.22 to 0.57, respectively) (**Figure 4A**). Although IgM antibodies were significantly correlated with complement (r = 0.64, p < 0.0001), the r values were notably higher for cytophilic IgG1 and IgG3 at 0.86 and 0.84, respectively (**Figure 4A**). Remarkably, GIA was not correlated with the different antibody isotypes and subclasses (**Figure 4A**).

**Figure 4:**
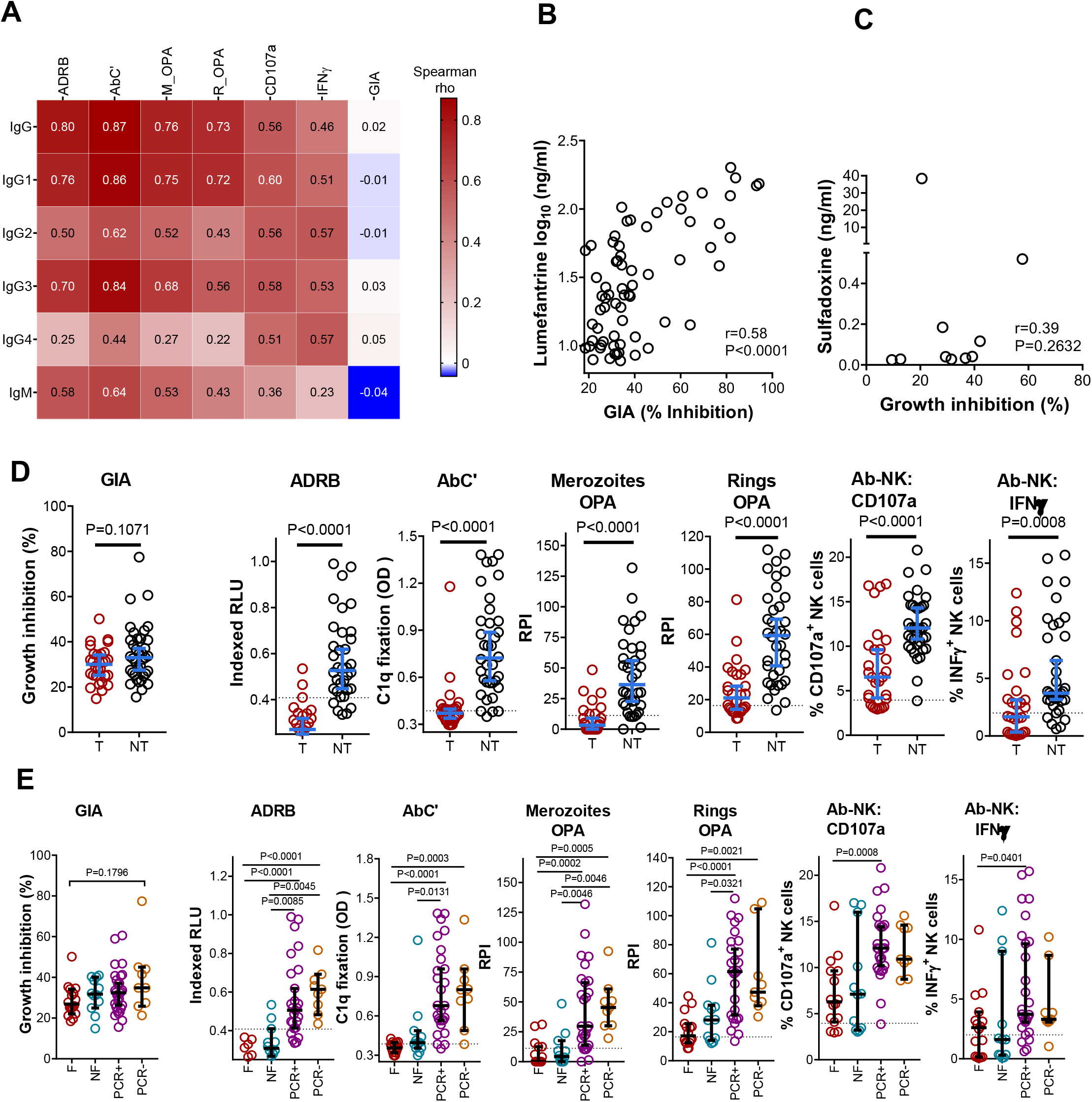
Fc-mediated effector functions were correlated with antibody levels but GIA was strongly associated with residual drug levels. **(A)** A heatmap showing Spearman’s correlation matrix of antibody effector functions and antibody levels. The colour intensity indicates the strength of the correlation. The Spearman’s rho value for each pair of antibodies is shown, n=142. All correlations were significant (*p* <0.05) except between GIA and all other variables, and between NK cell IFNγ production and IgM antibodies. **(B)** Spearman’s correlation for GIA and lumefantrine in volunteers with sub-therapeutic lumefantrine levels (n=68). **(C)** Spearman’s correlation for GIA and sulfadoxine in volunteers with sub-therapeutic sulfadoxine levels (n=10). **(D)** GIA (neutralization) and Fc-mediated effector functions were compared between treated (T, n=28) and non-treated (NT, n=36) in drug-negative volunteers. **(E)** GIA and Fc mediated antibody effector functions were compared across the four clinical endpoints, treated febrile (F, n=16), treated non-febrile (NF, n=12), not treated PCR positive (PCR+, n=27) and not treated PCR negative (PCR-, n=9) in drug-negative volunteers. Error bars represent median and 95% confidence intervals. P values were calculated using Mann-Whitney test for treatment outcome and using the Kruskal Wallis test with Dunn’s multiple comparisons test for the different clinical endpoints. The dotted black horizontal line represents the seropositivity cut-off (mean + 3SD of malaria naïve plasma samples). ADRB; antibody dependent respiratory burst by neutrophils, AbC’; complement fixation, OPA; opsonic phagocytosis by monocytes, ab-NK: Fc receptor-mediated natural killer (ab-NK) cell degranulation (CD107a) and IFNγ production, RLU; relative light units, RPI; relative phagocytosis index.

### Sub-therapeutic levels of antimalarial drugs confound analyses of immune correlates

We sought to better understand why antibodies inducing GIA were not correlated with anti-merozoite antibodies. We questioned whether the presence of residual antimalarial drugs in the plasma inhibited parasite growth in a similar manner to neutralizing antibodies in the GIA. Unexpectedly, sub-therapeutic levels of the antimalarial drugs lumefantrine and sulfadoxine (defined as below the minimum inhibitory concentrations) had been detected in retrospective analyses in 78/142 volunteers (n=68 for lumefantrine and n=10 for sulfadoxine; Kapulu *et al*.,2021, **Table S1**). Although we had deliberately dialyzed our plasma samples to remove any residual antimalarials before conducting the GIA (Persson *et al*., 2006), we observed a significant correlation between lumefantrine levels and GIA activity in the individuals found to have sub-therapeutic drug levels (lumefantrine: n = 68, *r* = 0.58, *p* < 0.0001, **Figure 4B**). This is consistent with a previous observation that lumefantrine can bind to high-density lipoproteins (HDLs) in plasma (Colussi *et al*., 1999). The HDLs form particles of ~10nm (Lund-Katz and Phillips, 2010) and were likely to have been retained in the 10 kDa dialysis columns we utilized, potentially accentuating growth inhibition. Although GIA was also weakly correlated with low sub-therapeutic levels of sulfadoxine in ten volunteers, this reached the threshold of positivity in only 3/10 samples (*r* = 0.39, *p* = 0.2632, **Figure 4C**). Notably, none of the six Fc-mediated effector functions were correlated with either drug (**Table S2**).

### Fc-mediated effector functions remain superior to GIA in drug-negative sub-group analysis

Although the drug levels detected retrospectively were below minimal inhibitory concentrations, we sought to minimize any potential confounding. Therefore, for all subsequent analyses, we excluded data from the 78 individuals in whom sub-therapeutic levels of antimalarial drugs were detected. Re-analysis of the GIA data showed no difference in GIA between treated and non-treated drug-negative volunteers (n = 64, **Figure 4D**) and across the four clinical immunity endpoints (**Figure 4E**). The initial and comparatively modest differences observed when samples from all volunteers had been included (**Figure 3A and 3B**) were lost. In contrast, all Fc-mediated effector functions remained significantly different between protected and susceptible drug-negative volunteers (n = 64, p <0.001, **Figure 4D**). Similar results were obtained in a receiver operating characteristic (ROC) curves analysis (AUC=0.62 for GIA versus 0.74-0.94 for Fc-mediated functions, **Figure S3**). Fc-mediated effector functions were also significantly higher in both PCR+ and PCR- volunteers than in those who were febrile in subgroup analyses (p < 0.01, **Figure 4E**). However, for Ab-NK, these subgroup differences were only significant in the PCR+ versus febrile comparison.

To further validate our results, we used a Cox proportional hazards model to reanalyse our data in the subset of drug-negative volunteers (n = 64). We dichotomized the data into high and low categories using statistically-derived thresholds (Murungi *et al*., 2013, Rono *et al*.,2013). All the Fc-mediated effector functions tested remained strongly associated with protection (Hazard ratios comparing high versus low levels of function were between 0.05 to 0.19, p <0.001, **Figure 5A**). In contrast, the GIA was not significantly associated with protection (HR 0.53, 95%CI (0.25-1.12), p = 0.095). The effect size for individual Fc-mediated effector functions was comparable and confidence intervals overlapped suggesting an equal contribution to protection.

**Figure 5:**
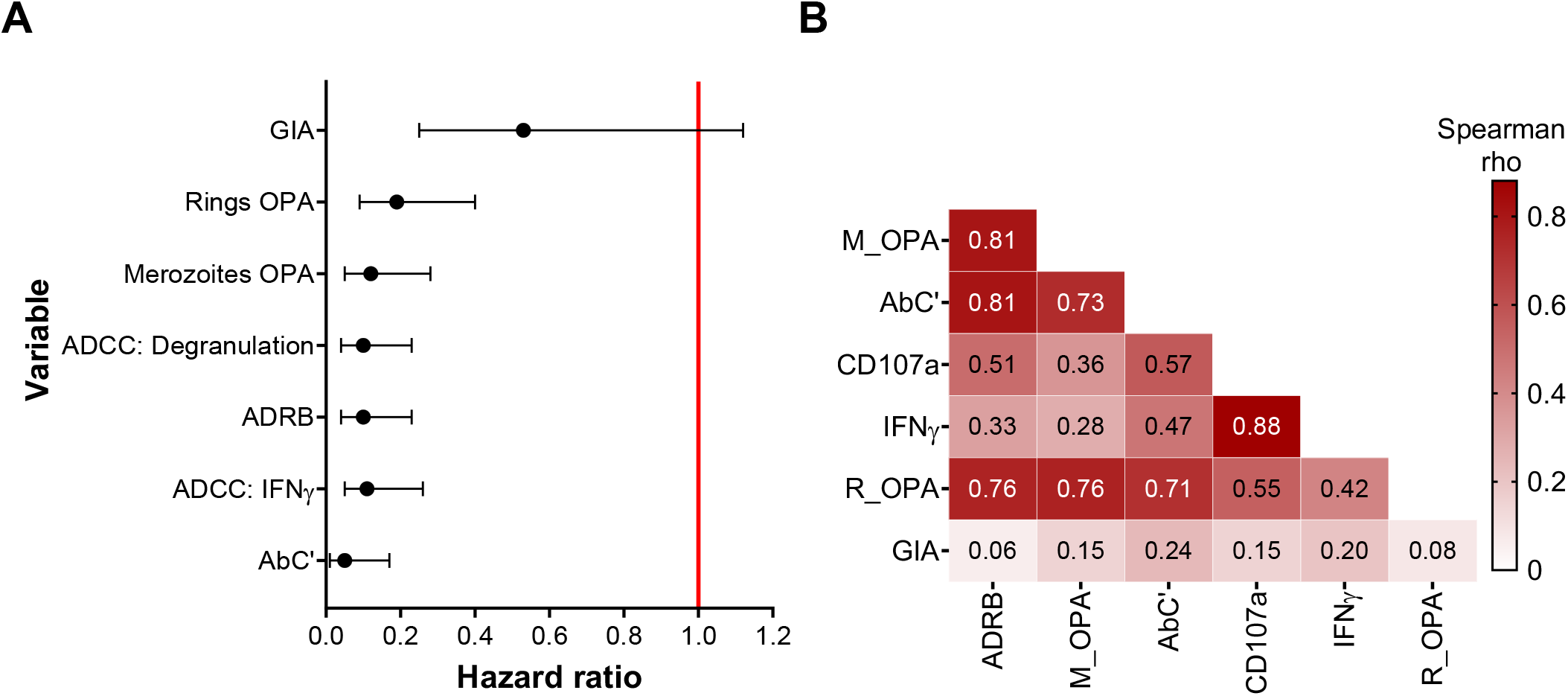
Fc-mediated effector functions were strongly associated with protection and intercorrelated. (**A**) A forest plot showing the adjusted Hazard ratios for each effector function using the Cox regression hazard model. The model compared the time to treatment between volunteers with responses high versus low levels of function. Error bars indicate 95% confidence intervals. The red line indicates no protection (Hazard ratio = 1.0). (**B**) A heatmap showing the Spearman’s correlation matrix for antibody effector functions. The colour intensity indicates the strength of the correlation. The Spearman’s rho values obtained for each pair of measures is shown. All correlations were significant (p <0.05) except between GIA and all other variablesADRB; antibody dependent respiratory burst by neutrophils, AbC’; complement fixation, M_OPA; opsonic phagocytosis of merozoites by monocytes, R_OPA; phagocytosis of ring stage parasites, CD107a; Fc-mediated NK cell degranulation, IFNγ; Fc-mediated NK cell IFNγ production. Data for drug-negative volunteers, n= 64.

We further tested whether Fc-mediated effector functions remained superior to neutralization when *in vivo* parasite densities were considered. We tested correlations between antibody effector functions and parasite densities post challenge. There was only a weak correlation between GIA and the maximum and mean parasite densities (**Table S3**). In contrast, there was a stronger negative correlation between Fc-mediated effector functions with mean and maximum parasitemia (**Table S4**). Taken together, these findings are consistent with previous studies that showed Fc-mediated effector functions are associated with the control of parasite density and protection from clinical symptoms of malaria in cohort studies in children (Joos *et al*., 2010; Hill *et al*., 2013; Osier *et al*., 2014a; Boyle *et al*., 2015; Tiendrebeogo *et al*., 2015; Feng *et al*., 2021)

### Fc-mediated effector functions are correlated with one another

Next, we asked whether the antibody effector functions were correlated. Respiratory burst, merozoites and rings phagocytosis, and complement fixation functions were positively correlated (*r* values > 0.70, p <0.0001, **Figure 5B)**. As expected, Ab-NK cell degranulation and IFNγ production were positively correlated (*r* = 0.88, *p* < 0.0001). We observed that Ab-NK activity was the least strongly positively correlated with other Fc-mediated effector functions. In contrast, GIA was not correlated with any of the Fc-mediated effector functions (*r* < 0.25, *p* > 0.05, **Figure 5B**). These data suggest that antibodies inducing Fc-mediated functions are co-acquired or polyfunctional.

### Breadth of antibody Fc-mediated effector function predicts complete protection against malaria

We undertook a principal component analysis (PCA) of the six Fc-mediated effector functions to visualize how well they discriminated our clinical immunity endpoints. This initial analysis did not discriminate protected from susceptible drug-negative volunteers (**Figure 6A**). The breadth of the antibody response has previously been shown to be an important correlate of protection (Osier *et al*., 2008; Osier *et al*., 2014b; Murungi *et al*., 2016; Obiero *et al*., 2019; Proietti *et al*., 2020). We hypothesized that this failure of the PCA to discriminate subgroups was due to variation in the breadth of Fc function, which was likely to be highest in the most immune (PCR-) group. To test this, we created a breath score defined as the sum of high-level responses to the six Fc-mediated functions and compared this across subgroups. In keeping with our hypothesis, the majority of PCR+ and PCR- volunteers had a breadth score of six while none of the febrile and non-febrile treated volunteers had responses to all six functions (**Figure 6B**). A middle group with a breadth score of between 3 and 5, likely accounted for the overlap in the PCA (**Figure 6A** and **Figure 6B**).

**Figure 6:**
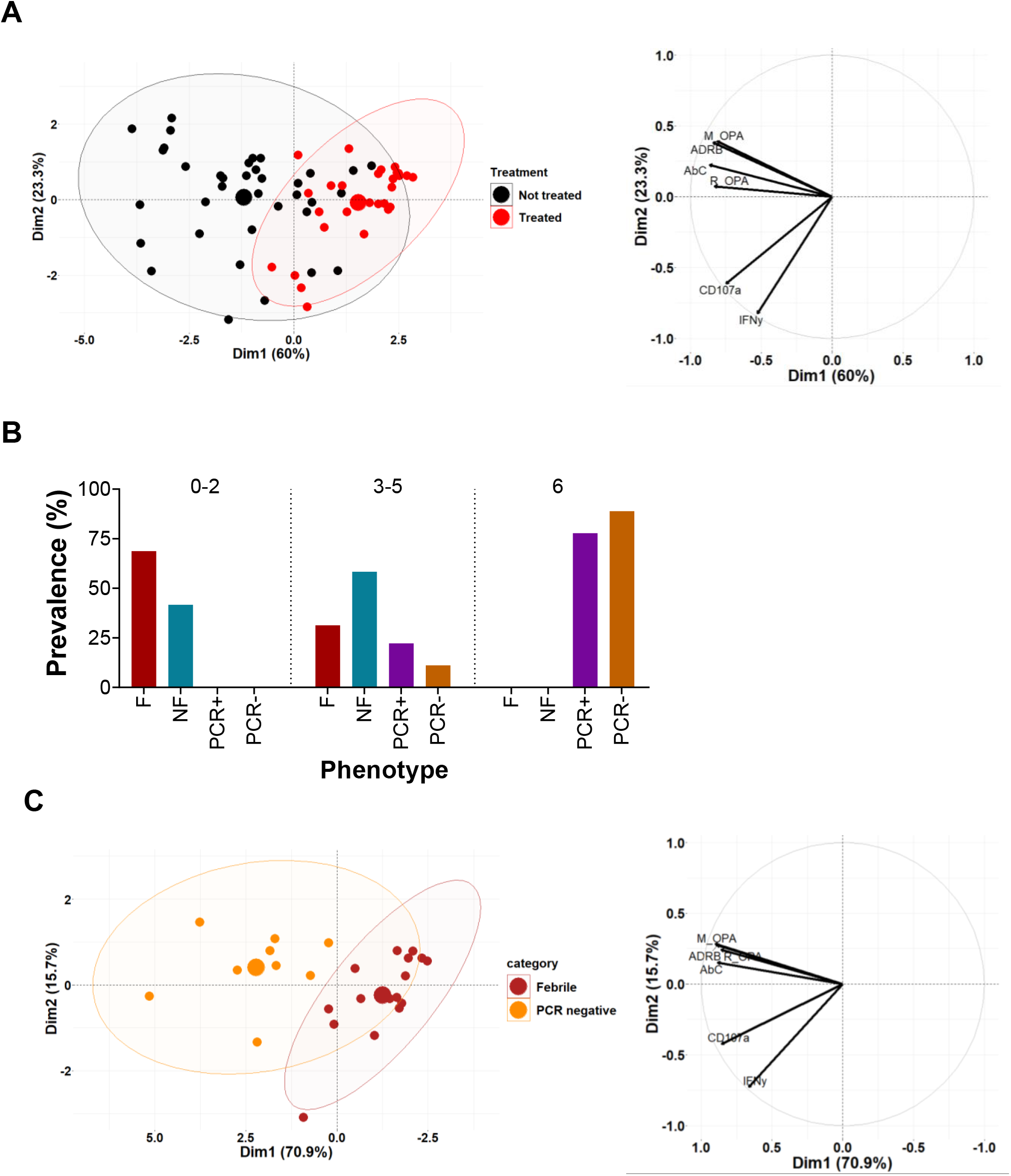
Overall protected volunteers had high levels Fc-mediated effector functions. (**A**) PCA for all the six Fc-mediated effector functions. The dots are coloured based on the treatment outcome, (treated; n=28, not treated; n=36). On the right is a loadings plot which mirrors the PCA dot plot indicating the location of the variables that affect the distribution of the individuals in the dot plot. (**B**) The proportion of volunteers with different breadth scores in each group. (**C**) PCA for all the six Fc-mediated effector functions. The dot plot coloured based on the two extreme endpoints (febrile; n=16 and PCR negative; n=9). On the right is a loadings plot which mirrors the PCA dot plot indicating the location of the variables that affect the distribution of the individuals in the dot plot. ADRB; antibody dependent respiratory burst by neutrophils, AbC’; complement fixation, M_OPA; opsonic phagocytosis of merozoites by monocytes, R_OPA; phagocytosis of ring stage parasites, CD107a: Fc-mediated NK cell granulation (CD107a), IFNγ: Fc-mediated NK cell IFNγ production. Data for drug-negative volunteers.

We therefore repeated a more stringent PCA comparing only the two extreme endpoints of clinical immunity (febrile versus PCR-) in the drug-negative sub-group. Our panel of six Fc-mediated effector functions fully discriminated these sub-groups (**Figure 6C**). We explored these data further and generated a heatmap to visualize the breadth of Fc function across the extremes of the clinical immunity endpoints. While it was immediately obvious that overall, febrile individuals had gaps in their Fc function repertoire, this was quite heterogenous with breadth scores ranging from 0/6 to 5/6 (**Figure 7A**). Likewise, 8/9 PCR-volunteers all had breadth scores of 6, while the 9^th^ had a score of 5/6 (**Figure 7A**). Additionally, 0/16 of the febrile volunteers had high-levels of complement mediated function, in contrast to 1/9 PCR-volunteers (**Figure 7A**).

**Figure 7:**
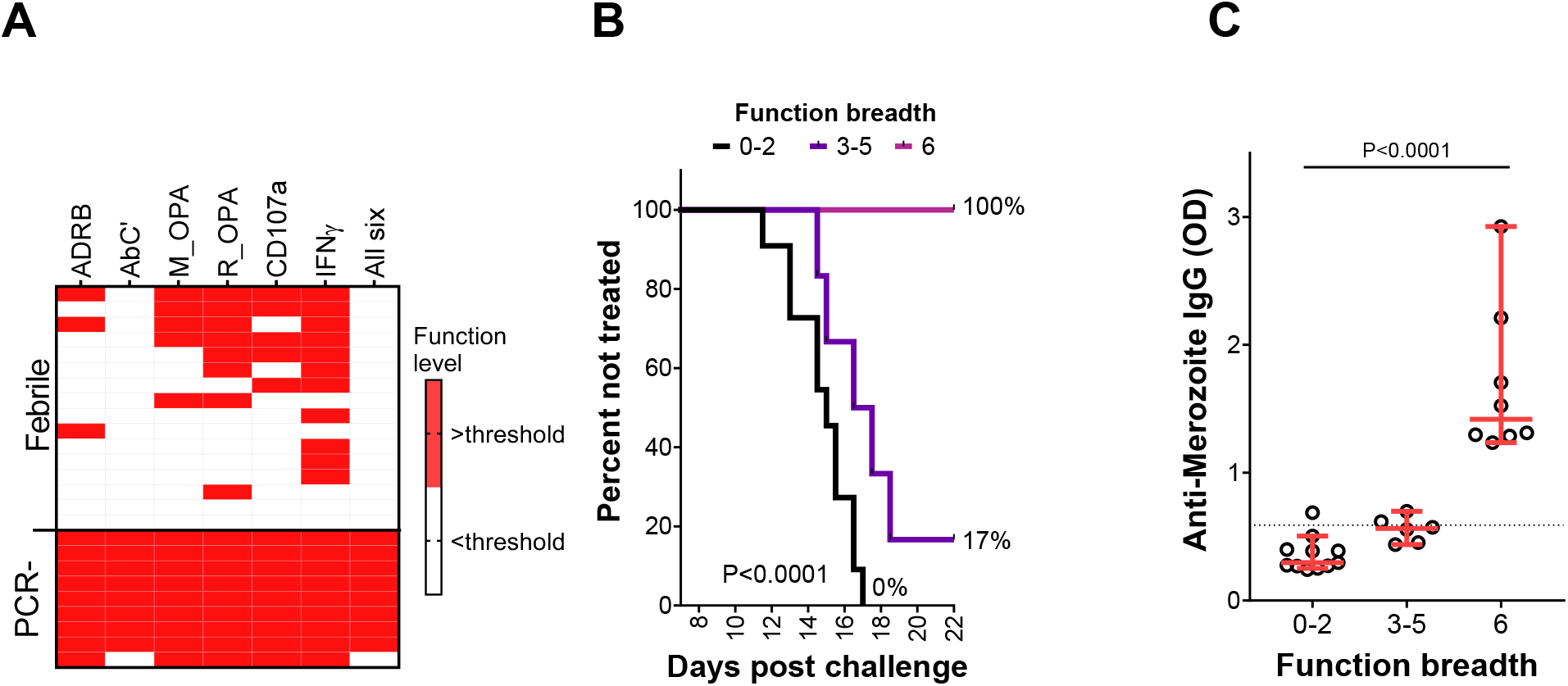
Breadth of antibody Fc-mediated effector function strongly predicts protection against malaria. (**A**) A heatmap of all six Fc-mediated effector functions in febrile (n=16) and PCR- (n=9) volunteers. High levels of function are highlighted in red. Each column represents an Fc-mediated function while each row represents a volunteer. ADRB; antibody dependent respiratory burst by neutrophils, AbC’; complement fixation, M_OPA; opsonic phagocytosis of merozoites by monocytes, R_OPA; phagocytosis of ring stage parasites, CD107a; Fc-mediated NK cell degranulation, IFNγ; Fc-mediated NK cell IFNγ production. (**B**) Survival curves showing the percentage of volunteers who remained untreated at different timepoints post challenge. Each line represents the breadth of function starting with 0-2 (n=11), 3-5 (n=6) and 6 (n=8). The p value was calculated using the Log-rank (Mantel-Cox) test. (**C**) Anti-merozoite IgG levels were compared between volunteers with varying breadth of Fc-mediated function. Each dot represents an individual. Error bars represent median and 95% confidence intervals. P values were calculated using Kruskal Wallis test. The dotted black horizontal line represents the seropositivity cut-off (mean + 3SD of malaria naïve plasma samples). See also Figure S3. Data for drug-negative volunteers.

We scrutinized these data further using Cox proportional hazards models. We observed a significant stepwise rise in protection with increasing breadth when comparing febrile versus PCR-ve volunteers. A breadth score of 0-2 Fc-mediated effector functions was not associated with protection, while a score of six was associated with complete protection (**Figure 7B**). A similar pattern was also observed when volunteers with other endpoints of clinical immunity (Non-febrile, PCR+) were included in the analysis (**Figure S4A & B)**. These data imply that the breadth of Fc-mediated effector functions is important for protection. Interestingly, the breadth of Fc-mediated effector functions increased with rising anti-merozoite IgG antibody levels (**Figure 7C; Figure S4C**). Taken together, the data suggests that the breadth of Fc-mediated effector functions is a better correlate of protection than any individual effector function.

To determine whether the differences in antibody levels observed between treated and non-treated volunteers and between different phenotypes were specific to *Plasmodium falciparum* antigens, we measured total IgG and total IgM antibodies. There was no significant difference in total antibodies between treated and non-treated volunteers (**Figure S5A & B**). Similarly, there was no significant difference across the four phenotype groups (**Figure S5A & B**) suggesting that the differences in antibody levels between volunteers with varied outcomes post CHMI are specific to antibodies against Plasmodium. However, the trends towards higher antibodies in the non-treated groups suggests that they generally respond better to a variety of antigens including those induced by malaria parasites.

## DISCUSSION

Although antibody mediated immunity to malaria has long focused on neutralization, an increasing body of evidence suggests that Fc-mediated effector functions should be prioritized as correlates of protection from malaria (Joos *et al*., 2010; Hill *et al*., 2013; Osier *et al*., 2014a; Boyle *et al*., 2015; Tiendrebeogo *et al*., 2015; Feng *et al*., 2021; Reiling *et al*., 2019). However, our understanding of the relative importance of different Fc-mediated effector functions in malaria is still limited as they have typically been analyzed in isolation, in unrelated studies. Here, we comprehensively analysed a panel of six antibody Fc-mediated effector functions and neutralization, and compared the strength of association between each mechanism and protection against experimental malaria in a single study.

Unlike previous study designs involving cohorts of children (Joos *et al*., 2010; Hill *et al*., 2013; Osier *et al*., 2014a; Boyle *et al*., 2015; Tiendrebeogo *et al*., 2015; Feng *et al*., 2021; Reiling *et al*., 2019), we used a CHMI approach that provides greater precision with regards to the timing, magnitude and quality of exposure (e.g. a single homogeneous parasite strain). Likewise, as volunteers were monitored in residential facilities, the clinical outcomes were captured as early as they occurred, documented accurately and enabled direct comparisons between volunteers. In line with previous studies, we found that Fc-mediated effector functions, but not neutralization, were strongly associated with protection from malaria. We further demonstrated that the breadth of Fc-mediated effector function was a stronger correlate of protection than any individual mechanism and enabled the clear discrimination of the extremes of clinical immunity following challenge.

We used the NF54 parasite strain that was isolated from one of the CHMI volunteers to minimize the possibility that differences between *in vitro* GIA activity and *in vivo* parasite growth were due to the parasite stain (Kennedy *et al*., 2002; Laurens *et al*.,2017). However, GIA was only very weakly correlated with parasitemia and was not significantly associated with protection post challenge. Antibodies against the *Pf*RH5 antigen have shown the highest GIA activity reported to date (Draper *et al*., 2018). *In vitro* GIA activity was correlated with *in vivo* parasite growth in Aotus monkeys immunized with *Pf*RH5 (Douglas *et al*., 2015). However, GIA inducing antibodies are required in prohibitively high concentrations for protection *in vivo*. Such concentrations have not yet been achieved in humans using current vaccination strategies. Indeed, the estimated amount of anti-*Pf*RH5 antibodies required for protection is very high, >300μg/ml (Douglas *et al*., 2015; Douglas *et al*., 2019). In contrast, naturally acquired antibodies against *Pf*RH5 were found in very low levels median <100ng/ml and max of 3μg/ml in adults from Kenya and Ghana (Payne *et al*., 2017). This suggests that these antibodies play a minor role in NAI.

Interestingly, GIA was correlated with plasma concentrations of antimalarial drugs, and was particularly evident for lumefantrine although the samples had been specifically dialysed prior to analysis to minimize this effect. Lumefantrine binds to high-density lipoproteins (Colussi *et al*., 1999) and likely persisted in plasma. Previous studies using samples from malaria exposed individuals that dialyzed samples did not measure lumefantrine levels (Dent *et al*.,2008; Murungi *et al*., 2016, 2017; Reiling *et al*., 2019). This may have confounded their results, especially in areas where artemisinin-lumefantrine is used as the first line of treatment for uncomplicated malaria (World Health Organization, 2015).

We found that high levels of Fc-mediated effector functions were characteristic of asymptomatic, low parasite density infections or complete parasite clearance whereas low levels of Fc-mediated functions were observed in febrile malaria. Taken together with earlier reports (Joos *et al*., 2010; Hill *et al*., 2013; Osier *et al*., 2014a; Boyle *et al*., 2015; Tiendrebeogo *et al*., 2015; Feng *et al*., 2021; Reiling *et al*., 2019), our findings suggest a decline in malaria symptoms and parasite density with increasing Fc-mediated antibody function. Unlike GIA, Fc-mediated effector functions remained significantly associated with protection even after excluding volunteers with detectable drug levels. Moreover, these Fc-mediated effector functions showed a moderate correlation with time to PCR positivity, time to treatment and *in vivo* parasite densities. Interestingly, this correlation was even stronger after excluding volunteers with detectable drug levels. These findings are consistent with studies that showed Fc-mediated effector functions were correlated with *in vivo* parasite growth (Bouharoun-Tayoun *et al*., 1990; Hodgson *et al*., 2016).

The effect size differed only marginally between the individual Fc-mediated effector functions suggesting that each contributes equally to protection. Polyfunctional antibodies capable of inducing multiple Fc-mediated effector functions may play an important role in parasite clearance and explain our findings of higher breadth of function in protected compared to susceptible volunteers. Indeed, the breadth of Fc-mediated effector functions clearly discriminated between volunteers with febrile malaria and those that were PCR negative. These findings support the clinical relevance of the breadth of Fc-mediated effector functions.

Unlike the GIA, all the Fc mediated effector functions were strongly correlated with anti-merozoite antibody levels in this study suggesting that this is likely their main mode of action. The breadth of Fc function was strongly correlated with antibody levels against merozoites. This suggests that variation in the expression of Fcγ receptors on the different effector cell types may contribute to the quality of the response (Lu *et al*., 2018), varied antibody concentrations may be required to activate the different receptors.

We and others have shown that invasion inhibition by anti-merozoite antibodies was significantly increased in the presence of NK cells (Odera *et al*., 2021) and complement (Boyle *et al*., 2015) suggesting that Fc-mediated effector mechanisms are induced rapidly and can target merozoites before invasion. Additionally, Boyle et al., 2015 showed that GIA activity was not associated with protection but growth inhibition in the presence of complement proteins was associated with protection. Similarly, certain neutralizing antibodies against Ebola (Gunn *et al*., 2018) influenza virus (DiLillo et al., 2014; Mullarkey et al., 2016), HIV (Hessell *et al*., 2007; Bournazos *et al*., 2014) and Hanta virus (Garrido *et al*., 2018) were more potent *in vivo* when in addition to neutralization they could also induce Fc-mediated effector functions. These studies suggest that although neutralizing antibodies are important in some infectious diseases, Fc-mediated functional activity is also highly relevant for protection. Consistent with previous studies, we observed that the prevalence and magnitude of naturally acquired IgG, mainly cytophilic subclasses, as well as IgM antibodies to merozoites was strongly correlated with protection (Bouharoun-Tayoun and Druilhe, 1992; Oeuvray *et al*.,2000; Stanisic *et al*., 2009; Boyle *et al*., 2019).

Malaria vaccine development has long prioritized vaccine candidates based on their ability to induce antibodies that inhibit parasite growth. However, antibodies mediate protection through a wide range of effector functions, many of which are Fc-mediated. Here we show that Fc-mediated effector functions were strongly associated with protection following experimental malaria challenge and that growth inhibition was not. Our data suggests that multiple Fc-mediated effector functions are likely necessary for protection against malaria and present a paradigm shift in our understanding of naturally acquired immunity that may impact the evaluation and prioritization of novel malaria vaccine candidates.

### Limitations

For safety reasons, the volunteers were treated at a parasitemia threshold of 500parasites/μl of blood regardless of whether or not they developed clinical symptoms of malaria. It is likely that at least a proportion of these individuals would have remained symptom free despite higher parasite densities and were thus misclassified as susceptible. Nonetheless, even with the conservative threshold of 500parasites/μl, we still observed marked differences in the time to treatment between volunteers.

We excluded volunteers with sickle cell trait as this is known to confer protection against malaria. However, other undetermined genetic factors may have contributed to the observed protection. Importantly, most genetic variants have a stronger impact on severe compared to mild malaria or parasitemia as measured in this study (Kariuki and Williams, 2020). Moreover, antibody responses to antigens expressed on the surface of infected red blood cells or sporozoites, as well as T cell responses, Fc receptor polymorphisms (Adu *et al*., 2012; Munde *et al*., 2017; Rohatgi *et al*., 2013), and donor specific differences in effector cell phenotypes and FcR expression (Hart *et al*., 2019, Damelang *et al*., 2019; Lu *et al*., 2018) may have contributed to the observed protection. Further studies will be needed to evaluate the impacts of each of these potential confounders.

Since the panel of assays tested was not exhaustive, it remains possible that additional Fc-mediated effector functions such as antibody dependent cellular inhibition (ADCI, Bouharoun-Tayoun *et al*., 1995), opsonic phagocytosis by neutrophils (Kumaratilake and Ferrante, 2000), and ADCC with γδ T cells (Farrington *et al*., 2020) contribute to protection. Nonetheless, our findings show that the breadth of the six-panel Fc-functions tested here clearly distinguished individuals demonstrating complete immunity from those succumbing to fever following a malaria challenge.

## Members of the CHMI-SIKA Study Team

Abdirahman I. Abdi^2^, Yonas Abebe^6^, Philip Bejon^2,7^, Peter F. Billingsley^6^, Peter C Bull^9^, Zaydah de Laurent^2^, Mainga Hamaluba^2^, Stephen L. Hoffman^6^, Eric R. James^6^, Melissa C. Kapulu^2^, Silvia Kariuki^2^, Domitila Kimani^2^, Rinter Kimathi^2^, Sam Kinyanjui^2,8,10^, Cheryl Kivisi^10^, Johnstone Makale^2^, Kevin Marsh^2,7^, Khadija Said Mohammed^2^, Moses Mosobo^2^, Janet Musembi^2^, Jennifer Musyoki^2^, Michelle Muthui^2^, Jedidah Mwacharo^2^, Kennedy Mwai^2,5^, Francis Ndungu^2^, Joyce M. Ngoi^2^, Patricia Njuguna^2^, Irene N. Nkumama^1,2^, Omar Ngoto^2^, Dennis O. Odera^1,2^, Bernhards Ogutu^8,11^, Fredrick Olewe^8^, Donwilliams Omuoyo^2^, John Ong’echa^8^, Faith H. A. Osier^1,2,5^, Edward Otieno^2^, Jimmy Shangala^2^, Betty Kim Lee Sim^6^, Thomas L. Richie^6^, James Tuju^2,4^, Juliana Wambua^2^, Thomas N Williams^2,12^.

### Affiliations

^6^Sanaria Inc., Rockville, MD, USA

^7^Centre for Tropical Medicine and Global Health, Nuffield Department of Medicine, University Oxford, Oxford, UK

^8^Centre for Clinical Research, Kenya Medical Research Institute, Kisumu, Kenya

^9^Department of Pathology, University of Cambridge, Cambridge, UK

^10^Pwani University, P. O. Box 195-80108, Kilifi, Kenya

^11^Center for Research in Therapeutic Sciences, Strathmore University, Nairobi, Kenya

^12^Department of Medicine, Imperial College London, London, UK

## ACKNOWLEDGMENTS

We are grateful to all the study volunteers who have participated in the CHMI-SIKA study. We are also very grateful to the study teams at the study sites in Kilifi and Ahero, the collaborating teams at Sanaria, the study investigators, and all the clinical and laboratory teams. The CHMI-SIKA study was supported by a Wellcome Trust grant (107499) and sponsored by the University of Oxford. This work was supported by a Sofja Kovalevskaja Award from the Alexander von Humboldt Foundation (3.2 - 1184811 - KEN - SKP) and an EDCTP Senior Fellowship (TMA 2015 SF1001) which is part of the EDCTP2 programme supported by the European Union awarded to F.H.A.O. F.K.M. was supported by a scholarship from the German Academic Exchange Service (DAAD), Funding Programme 57214224, ST-32-PKZ 91608705. K.M was supported by an NIHR Global Health Research Unit grant number 16/136/33; Tackling Infections to Benefit Africa (TIBA). K.M was also supported through the DELTAS Africa Initiative Grant No. 107754/Z/15/Z-DELTAS Africa SSACAB and from DELTAS Africa Initiative [DEL-15-003]. The DELTAS Africa Initiative is an independent funding scheme of the African Academy of Sciences (AAS)’s Alliance for Accelerating Excellence in Science in Africa (AESA) and supported by the New Partnership for Africa’s Development Planning and Coordinating Agency (NEPAD Agency) with funding from the Wellcome Trust [107769/Z/10/Z] and the UK government. The views expressed in this publication are those of the author(s) and not necessarily those of AAS, NEPAD Agency, Wellcome Trust or the UK government.

## AUTHOR CONTRIBUTION

F.H.A.O. and I.N.N. conceived the study and and wrote the paper with contributions from R.F., D.O, F.M., P.N., M.H., and M.K.. The experiments were designed and performed by I.N.N., D.O., F.M., L.N., L.M., J.T., K.F., M.R., and R.K.. I.N.N, K.M., D.O., and F.M. analyzed the results. I.N.N. and K.M prepared the figures. All authors read and approved the final version of the manuscript.

## STAR METHODS

### RESOURCE AVAILABILITY

#### Lead contact

Further information and requests for resources and reagents should be directed to and will be fulfilled by Lead Contact, Prof. Faith Osier

#### Materials availability

This study did not generate new unique reagents.

#### Data and code availability

The study protocol and outcomes are published (Kapulu et al., 2018; Kapulu *et al*., 2021). The other original data that support the findings of this study are available from the corresponding author upon reasonable request;

### EXPERIMENTAL MODEL AND SUBJECT DETAILS

#### CHMI volunteers

The Controlled Human Malaria Infection of Semi-Immune Kenyan Adults (CHMI-SIKA) study protocol was previously described in detail (Kapulu *et al*., 2018). Briefly, 161 healthy malaria exposed Kenyan volunteers aged 18-45 years were enrolled into the study after signing an informed consent. The volunteers were from different malaria transmission settings in Kenya. The study was conducted in 3 cohorts between 2016 and 2018. The participants were infected with Sanaria^®^ 3,200 aseptic, purified, cryopreserved NF54 sporozoites by direct venous injection (DVI). Blood stage parasitemia was monitored by quantitative polymerase chain reaction (qPCR) twice a day from day 7 to day 14 then once a day from day 15 to day 21 post infection (Kapulu *et al*., 2021). During the 21 day follow up period, volunteers were treated with artemether-lumefantrine (AL) if they had parasite densities above 500 parasites/μl of blood or if they had signs and symptoms of malaria and detectable parasites. At the end of the follow up on day 22 all volunteers were treated to clear any remaining parasites.

Plasma samples collected the day before challenge (C-1) were used in this study. Volunteers with antimalarial drug levels (Lumefantrine, pyrimethamine, sulfadoxine, chloroquine, artesunate or artemether) above the minimum inhibitory concentration on day seven post challenge were excluded from analysis (Kapulu *et al*., 2021). Additionally, volunteers with parasite clones other than the NF54 strain used for challenge, as determined by MSP2 genotyping (Kapulu *et al*., 2021), were also excluded. A total of 142 volunteers were included in the analysis of which 69% were male.

The CHMI-SIKA study was conducted at the Kenya Medical Research Institute (KEMRI)-Wellcome Trust Research Programme in Kilifi, Kenya with ethical approval from the KEMRI Scientific and Ethics Review Unit (KEMRI//SERU/CGMR-C/029/3190) and by the University of Oxford Tropical Research Ethics Committee (OxTREC 2-16). All participants gave written informed consent. The study was registered on ClinicalTrials.gov (NCT02739763), conducted based on good clinical practice (GCP), and under the principles of the Declaration of Helsinki.

#### Cell lines

The human monocyte cell line, THP-1 cells (sourced from ATCC), were cultured in RPMI 1640 medium supplemented with 2mM L-glutamine, 10 mM HEPES, 1% pen strep (10,000 units/ml penicillin and 10,000 μg/ml streptomycin) and 10% foetal bovine serum (FBS) in a humidified incubator with 5% CO_2_ at 37°C.

### METHOD DETAILS

#### Quantitative Polymerase Chain Reaction (qPCR)

The qPCR assay was described in detail elsewhere (Kapulu *et al*., 2021). Briefly, DNA was extracted from 500 μl of whole blood and eluted in 100 μl. An aliquot of 13.5 μl of the DNA was used for qPCR using a TaqMan^®^ probe (5’-FAM-AACAATTGGAGGGCAAG-NFQ-MGB-3’) which targets the multicopy 18S ribosomal RNA gene. Six serial dilutions of DNA extracted from an *in vitro* parasite culture of known parasitemia were included as standards and used to determine the parasitemia of the samples.

#### Merozoite isolation

*Plasmodium falciparum* merozoites of the NF54 or 3D7 strains were isolated as previously described (Boyle *et al*., 2010). Parasites were thawed and cultured to obtain trophozoites at high parasitemia (8-12%). The trophozoites were isolated by magnetic purification and cultured in fresh complete medium to allow development to early schizont stage. A protease inhibitor, trans-epoxysuccinyl-L-leucylamido (4-guanidino) butane (E64, Sigma Aldrich), was added to allow maturation of the schizonts without rapture (Boyle *et al*., 2010). Merozoites were harvested by filtration of the mature schizonts through 1.2μm filters (Pall). The merozoites were stained using either 1μg/ml ethidium bromide (EtBr, Thermo Fischer) or with 1X SYBR green dye (Thermo Fischer) for 30min and counted against CountBright™Absolute Counting Beads (Thermo Fischer) using BD FACS Canto II flow cytometer.

#### Anti-Merozoite ELISAs

96-well plates (Thermo Fischer) were coated with 100μl/well of NF54 merozoites at 5×10^6^ merozoites/ml at 4°C overnight. The plates were washed and blocked with 200μl/well of blocking buffer, 1% casein (Thermo Fischer), for 2 hours at 37°C. Samples were diluted at 1:500 with blocking buffer, added at 200μl/well and incubated for 1 hour at 37°C. Plates were washed before 100μl/well of respective secondary antibodies, Horseradish peroxidase (HRP)-conjugated: rabbit anti-human IgG (Dako), goat anti-human IgM (Southern Biotech) and rabbit anti-human IgG1, 2, 3 or 4 (The Binding Site GmbH), were added and incubated for 1 hour at 37°C. The plates were then washed four times and 100μl/well of substrate (o-phenylenediamine dihydrochloride (OPD), Sigma-Aldrich) was added and incubated for 20 min in the dark at room temperature (RT). The reaction was stopped with 30μl of 1M Hydrochloric acid (HCl, Sigma-Aldrich) and the absorbance measured at 492nM using BioTek Cytation 3 cell imaging multi-mode reader.

#### Tetanus toxoid ELISA

The protocol for tetanus toxoid ELISA is similar to the merozoite ELISA protocol above with a few variations. Briefly, plates were coated with 2μg/ml of tetanus toxoid (NIBSC), diluted in carbonate-bicarbonate buffer. Plasma samples were tested at 1:1000 dilution. Bound antibodies were detected with rabbit anti-human IgG conjugated HRP (Dako). The substrate (OPD) was added and incubated for 15 min in the dark at RT. The reaction was stopped with 30μl of 1M Hydrochloric acid (HCl) and the absorbance measured at 492nM.

#### Total IgG and IgM sandwich ELISAs

ELISA plates (Thermo Fischer) were coated overnight at 4°C with 10μg/ml of unlabelled rabbit anti-human IgG (Southern Biotech) or 2.5μg/ml of unlabelled goat anti-human IgM (Southern Biotech) diluted in bicarbonate buffer. The plates were blocked with 200μl of blocking buffer (3% skimmed milk) for 1 hour at RT. Plasma samples were diluted to 1:500,000 or 1:5000 for tIgG and tIgM ELISAs, respectively. Bound antibodies were detected with either HRP-conjugated rabbit anti-human IgG (Dako) or Alkaline phosphatase-conjugated goat antihuman IgM (Southern Biotech) for tIgG and tIgM ELISAs, respectively. The substrate (OPD or p-Nitrophenyl Phosphate and disodium Salt (PNPP) for tIgG and tIgM ELISAs, respectively) were added and incubated for 5-15 min in the dark at RT before the absorbance was read at 492nM or 405nm for tIgG and tIgM ELISAs, respectively.

#### Growth inhibition activity (GIA)

Plasma samples were dialysed using 10kDa columns (Merck Millipore GmbH), to remove drugs then heat inactivated at 56°C for 30min, to inactivate complement proteins. The samples were transferred into u-bottomed 96-well plates (Thermo Fischer) at 5μl/well. A few control wells with medium alone were also included as reference to calculate percentage inhibition. Tightly synchronized trophozoite stage parasites at 0.5% and 1% haematocrit were added into each well, 45μl/well. The parasites were incubated for 96 hours (2 cycles). At 48 hours, 10μl of fresh medium was added to each well. After 96 hours, the parasites were stained with 1x SYBR Green, washed and fixed in 200μl of 2% paraformaldehyde (PFA, AppliChem). The parasitemia in each well was then measured by flow cytometry using the high throughput sampler (HTS) on BD FACS Canto II™ flow cytometer. The FACs data was analysed in FlowJo version 10 software. Percentage parasitemia was determined as the percentage of SYBR green positive erythrocytes. Percentage inhibition was calculated as: 100 - (parasitemia of test sample / parasitemia of reference x 100).

#### Antibody dependent complement fixation (AbC’)

Complement fixation was measured using a previously described modified ELISA protocol (Boyle *et al*., 2015; Reiling *et al*., 2019). Briefly, 96-well ELISA plates (Thermo Fischer) were coated with 100μl of freshly isolated merozoites at 5×10^6^ merozoites/ml for 2 hours at 37°C. Plates were washed and blocked with 200μl of 1% Casein in PBS for 2 hours at 37°C. Plasma samples were diluted 1:100 in 0.1% casein in PBS, added to the plates 50μl/well and incubated for 2 hours at 37°C. The plates were then washed and 40μl of human C1q (Calbiochem) was added at 10μg/ml and incubated for 30 min at RT. The bound C1q was detected using HRP-conjugated sheep polyclonal anti-C1q antibody (Abcam) diluted 1:100 for 1 hour at 37°C. The plates were washed and 100μl of OPD substrate was added and incubated for 15 min at RT before the reaction was stopped by addition of 25μl of 2M H_2_SO_4_ and absorbance read at 492nm.

#### Opsonic phagocytosis activity (OPA) of merozoites

The OPA assay was done as previously described (Osier *et al*., 2014a). THP-1 cells were cultured and diluted to 6.7×10^5^ cells/ml. 96-well U-bottom plates were pre-coated with 1% casein in PBS for 20 min. THP-1 cells were transferred into the plates, 150μl/well, and placed in 37°C incubator. Plasma samples were diluted at 1:100 in RPMI, 10μl was transferred to the precoated plate and incubated with 100μl of ethidium bromide stained merozoites (8×10^6^merozoites/ml) for 40min at RT to allow opsonization to occur. For each sample, 50μl of opsonized merozoites were transferred in duplicates into the plates containing THP-1 cells and incubated at 37°C for 10 minutes to allow phagocytosis to occur.

Phagocytosis was stopped by centrifugation at 350 x g for 5 min at 4°C. Cells were washed once with 200μl ice cold FACS buffer (PBS + 0.5% BSA + 2mM EDTA) before they were resuspended in 200μl ice cold 2% PFA. Phagocytosis was measured using the high throughput sampler (HTS) on BD FACS Canto II™ flow cytometer. The data was analysed in FlowJo version 10 software. Phagocytosis was determined as the percentage of THP1-cells that were positive for ethidium bromide staining. A pool of hyperimmune serum from Kenyan adults (PHIS) was used as the reference sample. The relative phagocytosis index (RPI) of each sample was calculated as: phagocytosis of test sample / phagocytosis of reference sample (PHIS) X 100.

#### Opsonic phagocytosis activity (OPA) for ring stage parasites

The opsonic phagocytosis assay of ring stage parasites was previously described in detail (Musasia *et al*., 2022). Briefly, tightly synchronised ring stage parasites (0-10hours post infection) at 10-15% parasitemia were stained with 5μg/ml dihydroethidium (DHE, Abcam Limited) and μM Cell Trace™ violet dye (Thermo Fischer) in PBS for 30 min at 37°C. The stained ring culture was washed twice RPMI 1640 medium and transferred into u-bottomed 96-well plates at 0.5μl pellet per well. The ring culture was opsonised with heat-inactivated plasma diluted 1:12.5 for 30 minutes at RT. The opsonised cells were washed twice with RPMI 1640 and transferred in plates containing 150μl of THP1 cells (2.0 x 10^4^ cells/well) and incubated for 4 hrs at 37°C. Unphagocytosed erythrocytes were lysed. The cells were fixed with 2% PFA and phagocytosis was measured using the high throughput sampler (HTS) on BD FACS Canto II™ flow cytometer. The FACs data was analysed in the same manner as merozoite phagocytosis.

#### Antibody dependent respiratory burst (ADRB)

The ADRB assay was carried out as preciously described (Murungi *et al*., 2017). *Plasmodium falciparum* trophozoites of the 3D7 strain were purified by magnetic purification and then cultured in fresh complete medium to allow development to early schizont stage. A protease inhibitor E64 (Sigma Aldrich) was added to allow maturation of the schizonts for 8-12 hours without rapture. The schizonts were then counted and stored at −80°C before use. The schizonts were thawed and diluted to 10 x 10^5^ schizonts/ml in PBS and used to coat white opaque 96-well plates (Greiner), 100μl/well, overnight at RT. The plates were washed and blocked with 200μl/well casein in PBS for 1 hour at room temperature. Plasma samples were diluted 1:50 in 1X PBS. The plates were washed before 50μl of diluted plasma was added to each well and incubated at 37°C for 1 hour.

Polymorphonuclear leucocytes (PMN) were prepared fresh for each assay. Whole blood, 40ml, was collected from healthy donors and mixed at a ratio of 1:1 with Hank’s Balanced Salt Solution (HBSS, Thermo Fischer). The diluted blood was carefully layered on 7.5ml aliquots of Ficoll (GE healthcare). This was then centrifuged at 600xg for 15 min. The supernatant was carefully removed without disturbing the RBC pellet. The pellet was resuspended in 5ml of HBSS then mixed with 3% dextran at a ratio of 1:2 and incubated at room temperature in the dark for 30min. The supernatant was then carefully collected and centrifuged at 500 x g for 7min at 4°C. The supernatant was discarded, and RBC contaminants lysed. The cells were then centrifuged at 500 x g for 7min at 4°C and the pellet resuspended in 1ml of ice cold PMN buffer (HBSS with 0.1% bovine serum albumin (BSA, Sigma-Aldrich), 1% D -glucose). The PMN count determined using a haemocytometer and the concentration adjusted using PMN buffer to 1.0 × 10^7^ / ml. The cells were kept on ice.

The plates were washed and 50μl of 0.04mg/ml isoluminol (Santa Cruz Biotechnology) and 50μl of PMN were added to each well. The plates were immediately loaded onto the plate reader and ADRB activity measured as luminescence at 450nM every two minutes for 1.5 hours. The maximal relative light unit (RLU) values were obtained for analysis. Due to donor-donor variation in ADRB activity, each sample was run with PMNs from two different donors. The RLU values were indexed based on the positive control (PHIS) in each plate. The mean indexed RLU values using PMN from two donors was then calculated.

#### Antibody-mediated natural killer cells activation (Ab-NK)

The antibody-mediated natural killer cells activation (Ab-NK) assay has been previously described (Odera *et al*., 2021). Briefly, 96 well culture-plates were coated with 100μl of merozoites in PBS overnight at 4°C. The next day, the plates were washed and blocked with casein in PBS for 2 hours at 37°C before 50μl/well of plasma samples diluted 1:20 in blocking buffer were added and incubated for 4 hours at 37°C. Peripheral blood mononuclear cells (PBMCs) were harvested by density gradient centrifugation from fresh blood collected from healthy donors, resuspended in 2 ml of culture medium and the cell density determined using a hemocytometer. NK cells were enriched from the PBMCs using a NK negative isolation kit (Miltenyi) and magnetic cell sorting using LS columns (Miltenyi) as per the manufacture’s instructions and stored at 1X10^5^ cells per ml. Freshly isolated NK cells (20,000 NK cells/well), 2μl/well of brefeldin A (BFA, Sigma-Aldrich), 2μl/well of monensin (Sigma) and 2.5μ/well of PE conjugated anti-human CD107a antibody (BD biosciences) was added into the opsonised merozoites in the plates and incubated for 18 hours at 4°C. The next day, the stimulated NK cells were transferred into V bottom 96 well plates, centrifuged at 800 x g for 5 min at 4°C and washed with FACS buffer (0.1% Sodium Azide + 1% BSA + 1 x PBS). Cells were stained with 10μl of viability dye Live/dead FITC stain (BD biosciences) incubated at 4°C for 10min. NK cell surface receptors were stained with 20μl/well of: anti-human CD3-PE-Cy5, anti-human CD56-APC and anti-human CD16-APC-Cy7 (all from BD biosciences) for 30 min at 4°C. The plates were then washed 2 times with FACS buffer before the cells were permeabilized with 80μl of permeabilization buffer (permwash, BD biosciences) for 10 min at 4°C. The cells were stained with anti-human IFNγ-PE-Cy7 antibody (BD biosciences) diluted 1:50 in permwash and incubated for 1 hour at 4°C. The stained cells were washed with permwash and resuspended in 150μl of FACs buffer and stored at 4°C awaiting acquisition on BD FACS Canto II™ flow cytometer. Positive and negative control beads were included for each antibody and used as compensation controls. The proportion of NK cells expressing IFNγ and or CD107a (degranulating NK cells) were determined using FlowJo version 10.

### QUANTIFICATION AND STATISTICAL ANALYSIS

Statistical tests are indicated in the corresponding figure legends. Dot plots represent median with 95% confidence interval. A significance threshold of p < 0.05 was used for all tests.

#### Mann-Whitney U test

The nonparametric Mann-Whitney U test was performed using Graphpad Prism to compare antibody responses between different treated and non-treated volunteers.

#### Kruskal-Wallis H test with Dunn’s multiple comparisons test

Kruskal-Wallis H test with Dunn’s multiple comparisons test was performed using Graphpad Prism to compare antibody responses across the four phenotypes (febrile, nonfebrile, PCR positive and PCR negative volunteers) and in volunteers with different Fc-function breadths.

#### Receiver operating characteristic (ROC) curves

The performance of the different effector functions was evaluated using area under the receiver operating characteristic (ROC) curves. For each ROC curve, the area under the curve (AUC) is reported.

#### Correlation analysis

Spearman’s rank correlation was used to examine correlations between variables.

#### Threshold analysis

The antibody/function specific thresholds were determined using maximally selected rank statistics analysis method in R, as the cut-off of antibody/function levels with the most significant relation with time to treatment (Murungi *et al*., 2013, Rono *et al*., 2013).

#### Fc-mediated function breadth analysis

Fc-mediated effector functions breadth score was calculated in STATA by assigning a score of 1 to levels above threshold of each function and calculating the total score for each volunteer (Osier *et al*., 2008; Osier *et al*., 2014a; Osier *et al*., 2014b).

#### Survival analysis

Cox proportional hazards model was used to determine associations of antibody response or Fc-mediated effector function breadth with time to treatment adjusting for potential confounders (cohort and anti-malarial drug levels) in STATA. Kaplan-Meier curve with Log rank sum test in GraphPad Prism was used to compare survival curves of volunteers with different Fc-mediated function breadth.

#### Principal component analysis

Principal component analysis (PCA) was done using R to show Fc-mediated antibody effector functions that account for the highest variation between CHMI volunteers.

## SUPPLEMENTAL INFORMATION TITLES AND LEGENDS

**Figure S1: Protection was associated with high levels of antibodies against merozoites**

(**A**) The prevalence of IgG, IgM, and IgG 1-4 antibodies against merozoites and total IgG against tetanus toxoid antibodies compared across the four phenotypes based on parasite growth patterns, treated febrile (F, n=26), treated non-febrile (NF, n=30), not treated PCR positive (PCR+, n=53) and not treated PCR negative (PCR-, n=33). (**B**) IgG, IgM, and IgG 1-4 antibody levels against merozoites compared across the four phenotypes based on parasite growth patterns. (**C**) Total IgG against tetanus toxoid were compared between treated (n=56) and not treated (n=86) and across the four phenotypes based on parasite growth patterns. Error bars represent median and 95% confidence intervals. P values were calculated using Mann-Whitney test for treatment outcome and using Kruskal Wallis test with Dunn’s multiple comparisons test for the different phenotypes. The dotted black horizontal line represents the seropositivity cut-off (mean + 3SD of malaria naïve plasma samples).

**Supplementary Figure S2: Fc-mediated effector functions were more strongly associated with protection than GIA**.

Receiver operating characteristic (ROC) curves for effector functions for all volunteers (n=142). The area under the ROC curve for each function is shown in the figure legend. ADRB; antibody dependent respiratory burst by neutrophils, AbC’; complement fixation activity, GIA; growth inhibition assay, Merozoite_OPA; opsonic phagocytosis of merozoites activity by monocytes, Rings_OPA; phagocytosis of ring stage parasites, Ab-NK_CD107a: antibody dependent granulation (CD107a) by natural killer cells, Ab-NK_IFNγ: antibody dependent IFNγ production by natural killer cells.

**Supplementary Table S1: Baseline characteristics of the CHMI-SIKA volunteers**

^a^ The following antimalarial drugs were measured in plasma samples collected on day 7 post challenge: Lumefantrine, pyrimethamine, sulfadoxine, chloroquine, artesunate and artemether. Of the 142 volunteers, 78 had detectable Lumefantrine or Sulfadoxine or both at levels below the minimum inhibitory concentrations (MIC, lumefantrine 200ng/ml, sulfadoxine 100ng/ml).

**Supplementary Table S2: Correlation of Fc-mediated effector functions with detectable lumefantrine and sulfadoxine drug levels**

ADRB; antibody dependent respiratory burst by neutrophils, AbC’; complement fixation activity, GIA; growth inhibition assay, Merozoite_OPA; opsonic phagocytosis of merozoites activity by monocytes, Rings_OPA; phagocytosis of ring stage parasites, Ab-NK_CD107a: antibody dependent granulation (CD107a) by natural killer cells, Ab-NK_IFNγ: antibody dependent IFNγ production by natural killer cells.

**Supplementary Figure S3: Fc-mediated effector functions were more strongly associated with protection than GIA**. Receiver operating characteristic (ROC) curves for effector functions for only drug-negative volunteers (n=64). The area under the ROC curve for each function is shown in the figure legend. ADRB; antibody dependent respiratory burst by neutrophils, AbC’; complement fixation activity, GIA; growth inhibition assay, Merozoite_OPA; opsonic phagocytosis of merozoites activity by monocytes, Rings_OPA; phagocytosis of ring stage parasites, Ab-NK_CD107a: antibody dependent granulation (CD107a) by natural killer cells, Ab-NK_IFNγ: antibody dependent IFNγ production by natural killer cells.

**Supplementary Table S3: Correlation of GIA with secondary outcome variables in drug negative samples** The mean parasite density was calculated is a geometric mean of parasite density observed between days 8.5 and 22 after challenge, excluding timepoints after treatment; the maximum parasite density is the highest parasite density observed between days 8.5 and 22 after challenge, excluding timepoints after treatment. Data for drug-negative volunteers, n=64.

**Supplementary Table S4: Correlation of anti-merozoite Fc mediated effector functions with secondary outcome variables in drug negative samples** Spearman’s correlation. The mean parasite density was calculated is a geometric mean of parasite density observed between days 8.5 and 22 after challenge, excluding timepoints after treatment; the maximum parasite density is the highest parasite density observed between days 8.5 and 22 after challenge, excluding timepoints after treatment. ADRB; antibody dependent respiratory burst by neutrophils, AbC’; complement fixation activity, OPA; opsonic phagocytosis activity by monocytes, Ab-NK_CD107a: antibody dependent granulation (CD107a) by natural killer cells, Ab-NK_IFNγ: antibody dependent IFNγ production by natural killer cells. Data for drug-negative volunteers, n= 64.

**Supplementary Figure S4: The breadth of Fc-mediated effector functions was more strongly associated with protection than any individual function.**

(**A**) A heatmap of all six Fc mediated effector functions in treated (n=28) and non treated (n=36) volunteers. Responses above function specific thresholds derived using maximally selected rank statistics are highlighted in red. Each column represents an Fc-mediated function while each row represents a volunteer. ADRB; antibody dependent respiratory burst by neutrophils, AbC’; complement fixation activity, M_OPA; opsonic phagocytosis of merozoites activity by monocytes, R_OPA; phagocytosis of ring stage parasites, Degran: antibody dependent granulation (CD107a) by natural killer cells, IFNγ: antibody dependent IFNγ production by natural killer cells. (**B**) Survival curves showing percentage of volunteers who remained untreated at different timepoints post challenge. Each line represents a function breath level starting with 0-2 (n=16), 3-5 (n=19) and 6 (n=29). The p value was calculated using Log-rank (Mantel-Cox) test. (**D**) Anti-merozoite IgG levels were compared between volunteers with varying breadth of Fc mediated function. Each dot represents an individual. Error bars represent median and 95% confidence intervals. P values were calculated using Kruskal Wallis test. The dotted black horizontal line represents the seropositivity cut-off (mean + 3SD of malaria naïve plasma samples). Data for drug-negative volunteers, n = 64.

**Supplementary Figure 5: No significant differences in total antibody levels between protected and susceptible.**

(**A**)Total IgG and (**B**) total IgM antibodies were compared between volunteers who required treatment post challenge (n=28) and those who did not (n=36) and across the four phenotypes based on parasite growth patterns (Febrile n=16, non-febrile n=12, PCR positive n=27 and PCR negative n=9). Error bars represent median and 95% confidence intervals. p values were calculated using Mann-Whitney test for treatment outcome and using Kruskal Wallis test with Dunn’s multiple comparisons test for the different phenotypes. Data for drug-negative volunteers, n = 64.

## REFERENCES

Adu, B. et al. (2012) ‘Fc Gamma Receptor IIIB (FcγRIIIB) Polymorphisms Are Associated with Clinical Malaria in Ghanaian Children’, PLOS ONE. Public Library of Science, 7(9), p. e46197.

Bouharoun-Tayoun, H. et al. (1990) ‘Antibodies that protect humans against Plasmodium falciparum blood stages do not on their own inhibit parasite growth and invasion in vitro, but act in cooperation with monocytes.’, The Journal of experimental medicine. UNITED STATES, 172(6), pp. 1633–1641.

Bouharoun-Tayoun, H. et al. (1995) ‘Mechanisms underlying the monocyte-mediated antibody-dependent killing of Plasmodium falciparum asexual blood stages.’, The Journal of Experimental Medicine, 182(2), pp. 409–418. doi: 10.1084/jem.182.2.409.

Bouharoun-Tayoun, H. and Druilhe, P. (1992) ‘Antibodies in falciparum malaria: what matters most, quantity or quality?’, Memorias do Instituto Oswaldo Cruz. BRAZIL, 87 Suppl 3, pp. 229–234.

Bournazos, S. et al. (2014) ‘Broadly neutralizing anti-HIV-1 antibodies require Fc effector functions for in vivo activity.’, Cell, 158(6), pp. 1243–1253. doi: 10.1016/j.cell.2014.08.023.

Boyle, M. J. et al. (2010) ‘Isolation of viable Plasmodium falciparum merozoites to define erythrocyte invasion events and advance vaccine and drug development’. doi: 10.1073/pnas.1009198107/-/DCSupplemental.www.pnas.org/cgi/doi/10.1073/pnas.1009198107.

Boyle, M. J. et al. (2015) ‘Human antibodies fix complement to inhibit Plasmodium falciparum invasion of erythrocytes and are associated with protection against malaria.’, Immunity. United States, 42(3), pp. 580–590. doi: 10.1016/j.immuni.2015.02.012.

Boyle, M. J. et al. (2019) ‘IgM in human immunity to Plasmodium falciparum malaria.’, Science advances, 5(9), p. eaax4489. doi: 10.1126/sciadv.aax4489.

Colussi, D. et al. (1999) ‘Binding of artemether and lumefantrine to plasma proteins and erythrocytes.’, European journal of pharmaceutical sciences: official journal of the European Federation for Pharmaceutical Sciences. Netherlands, 9(1), pp. 9–16. doi: 10.1016/s0928-0987(99)00037-8.

Damelang, T. et al. (2019) ‘Role of IgG3 in Infectious Diseases.’, Trends in immunology. England, 40(3), pp. 197–211. doi: 10.1016/j.it.2019.01.005.

Dent, A. E. et al. (2008) ‘Antibody-mediated growth inhibition of Plasmodium falciparum: relationship to age and protection from parasitemia in Kenyan children and adults.’, PloS one, 3(10), p. e3557. doi: 10.1371/journal.pone.0003557.

DiLillo, D. J. et al. (2014) ‘Broadly neutralizing hemagglutinin stalk-specific antibodies require FcγR interactions for protection against influenza virus in vivo.’, Nature medicine, 20(2), pp. 143–151. doi: 10.1038/nm.3443.

Douglas, A. D. et al. (2015) ‘A PfRH5-based vaccine is efficacious against heterologous strain blood-stage Plasmodium falciparum infection in aotus monkeys.’, Cell host & microbe, 17(1), pp. 130–139. doi: 10.1016/j.chom.2014.11.017.

Douglas, A. D. et al. (2019) ‘A defined mechanistic correlate of protection against Plasmodium falciparum malaria in non-human primates’, Nature Communications, 10(1), p. 1953. doi: 10.1038/s41467-019-09894-4.

Draper, S. J. et al. (2018) ‘Malaria Vaccines: Recent Advances and New Horizons.’, Cell host & microbe, 24(1), pp. 43–56. doi: 10.1016/j.chom.2018.06.008.

Duncan, C. J. A., Hill, A. V. S. and Ellis, R. D. (2012) ‘Can growth inhibition assays (GIA) predict blood-stage malaria vaccine efficacy?’, Human vaccines & immunotherapeutics. United States, 8(6), pp. 706–714. doi: 10.4161/hv.19712.

Farrington, L. A. et al. (2020) ‘Opsonized antigen activates Vδ2+ T cells via CD16/FCγRIIIa in individuals with chronic malaria exposure’, PLOS Pathogens. Public Library of Science, 16(10), p. e1008997.

Feng, G. et al. (2021) ‘Mechanisms and targets of Fcγ-receptor mediated immunity to malaria sporozoites.’, Nature communications, 12(1), p. 1742. doi: 10.1038/s41467-021-21998-4.

Forthal, D. N. (2014) ‘Functions of Antibodies.’, Microbiology spectrum. United States, 2(4), p. AID-0019-2014. doi: 10.1128/microbiolspec.AID-0019-2014.

Garrido, J. L. et al. (2018) ‘Two recombinant human monoclonal antibodies that protect against lethal Andes hantavirus infection in vivo.’, Science translational medicine. United States, 10(468). doi: 10.1126/scitranslmed.aat6420.

Gunn, B. M. et al. (2018) ‘A Role for Fc Function in Therapeutic Monoclonal Antibody-Mediated Protection against Ebola Virus.’, Cell host & microbe, 24(2), pp. 221–233.e5. doi: 10.1016/j.chom.2018.07.009.

Hessell, A. J. et al. (2007) ‘Fc receptor but not complement binding is important in antibody protection against HIV.’, Nature. England, 449(7158), pp. 101–104. doi: 10.1038/nature06106.

Hill, D. L. et al. (2013) ‘Opsonising antibodies to P. falciparum merozoites associated with immunity to clinical malaria.’, PloS one, 8(9), p. e74627. doi: 10.1371/journal.pone.0074627.

Hodgson, S. H. et al. (2016) ‘Changes in serological immunology measures in UK and Kenyan adults post-controlled human malaria infection’, Frontiers in Microbiology. doi: 10.3389/fmicb.2016.01604.

Joos, C. et al. (2010) ‘Clinical protection from falciparum malaria correlates with neutrophil respiratory bursts induced by merozoites opsonized with human serum antibodies.’, PloS one, 5(3), p. e9871. doi: 10.1371/journal.pone.0009871.

Kapulu, M. C. et al. (2021) ‘Safety and PCR monitoring in 161 semi-immune Kenyan adults following controlled human malaria infection.’, JCI insight, 6(17). doi: 10.1172/jci.insight.146443.

Kapulu, M. C. et al. (2022) ‘Controlled human malaria infection (CHMI) outcomes in Kenyan adults is associated with prior history of malaria exposure and anti-schizont antibody response.’, BMC infectious diseases, 22(1), p. 86. doi: 10.1186/s12879-022-07044-8.

Kapulu, M. C., Njuguna, P. and Hamaluba, M. M. (2018) ‘Controlled Human Malaria Infection in Semi-Immune Kenyan Adults (CHMI-SIKA): a study protocol to investigate in vivo Plasmodium falciparum malaria parasite growth in the context of pre-existing immunity.’, Wellcome open research, 3, p. 155. doi: 10.12688/wellcomeopenres.14909.2.

Kariuki, S. N. and Williams, T. N. (2020) ‘Human genetics and malaria resistance’, Human Genetics, 139(6), pp. 801–811. doi: 10.1007/s00439-020-02142-6.

Kennedy, M. C. et al. (2002) ‘In vitro studies with recombinant Plasmodium falciparum apical membrane antigen 1 (AMA1): production and activity of an AMA1 vaccine and generation of a multiallelic response.’, Infection and immunity, 70(12), pp. 6948–6960. doi: 10.1128/iai.70.12.6948-6960.2002.

Kinyanjui, S. M. et al. (2009) ‘What you see is not what you get: implications of the brevity of antibody responses to malaria antigens and transmission heterogeneity in longitudinal studies of malaria immunity.’, Malaria journal. England, 8, p. 242. doi: 10.1186/1475-2875-8-242.

Kumaratilake, L. M. and Ferrante, A. (2000) ‘Opsonization and phagocytosis of Plasmodium falciparum merozoites measured by flow cytometry.’, Clinical and diagnostic laboratory immunology. UNITED STATES, 7(1), pp. 9–13.

Laurens, M. B. et al. (2017) ‘Strain-specific Plasmodium falciparum growth inhibition among Malian children immunized with a blood-stage malaria vaccine’, PLoS ONE. doi: 10.1371/journal.pone.0173294.

Lu, L. L. et al. (2018) ‘Beyond binding: antibody effector functions in infectious diseases.’, Nature reviews. Immunology, 18(1), pp. 46–61. doi: 10.1038/nri.2017.106.

Lund-Katz, S. and Phillips, M. C. (2010) ‘High density lipoprotein structure-function and role in reverse cholesterol transport.’, Sub-cellular biochemistry, 51, pp. 183–227. doi: 10.1007/978-90-481-8622-8_7.

Minassian, A. M. et al. (2021) ‘Reduced blood-stage malaria growth and immune correlates in humans following RH5 vaccination.’, Med (New York, N.Y.), 2(6), pp. 701–719.e19. doi: 10.1016/j.medj.2021.03.014.

Moormann, A. M., Nixon, C. E. and Forconi, C. S. (2019) ‘Immune effector mechanisms in malaria: An update focusing on human immunity.’, Parasite immunology. England, 41(8), p. e12628. doi: 10.1111/pim.12628.

Mullarkey, C. E. et al. (2016) ‘Broadly Neutralizing Hemagglutinin Stalk-Specific Antibodies Induce Potent Phagocytosis of Immune Complexes by Neutrophils in an Fc-Dependent Manner’, mBio. Edited by D. E. R. Griffin Guus Russell, Charles, 7(5), pp. e01624–16. doi: 10.1128/mBio.01624-16.

Munde, E. O. et al. (2017) ‘Association between Fcγ receptor IIA, IIIA and IIIB genetic polymorphisms and susceptibility to severe malaria anemia in children in western Kenya.’, BMC infectious diseases, 17(1), p. 289. doi: 10.1186/s12879-017-2390-0.

Murungi, L. M. et al. (2013) ‘A threshold concentration of anti-merozoite antibodies is required for protection from clinical episodes of malaria.’, Vaccine. Elsevier Ltd, 31(37), pp. 3936–42. doi: 10.1016/j.vaccine.2013.06.042.

Murungi, L. M. et al. (2016) ‘Targets and mechanisms associated with protection from severe Plasmodium falciparum malaria in Kenyan children’, Infection and Immunity. doi: 10.1128/IAI.01120-15.

Murungi, L. M. et al. (2017) ‘Cord blood IgG and the risk of severe Plasmodium falciparum malaria in the first year of life’, International Journal for Parasitology. doi: 10.1016/j.ijpara.2016.09.005.

Musasia, F. K. et al. (2022) ‘Phagocytosis of Plasmodium falciparum ring-stage parasites predicts protection against malaria’, Nature communications, Jul 14;13(. doi: 10.1038/s41467-022-31640-6.

Obiero, J. M. et al. (2019) ‘Antibody Biomarkers Associated with Sterile Protection Induced by Controlled Human Malaria Infection under Chloroquine Prophylaxis.’, mSphere, 4(1). doi: 10.1128/mSphereDirect.00027-19.

Odera, D. O. et al. (2021) ‘Antibodies targeting merozoites induce natural killer cell degranulation and interferon gamma secretion and are associated with immunity against malaria’, medRxiv, p. 2021.12.14.21267763. doi: 10.1101/2021.12.14.21267763.

Oeuvray, C. et al. (2000) ‘Cytophilic immunoglobulin responses to Plasmodium falciparum glutamate-rich protein are correlated with protection against clinical malaria in Dielmo, Senegal’, Infection and Immunity. doi: 10.1128/IAI.68.5.2617-2620.2000.

Osier, F. H. A. et al. (2008) ‘Breadth and magnitude of antibody responses to multiple Plasmodium falciparum merozoite antigens are associated with protection from clinical malaria.’, Infection and immunity, 76(5), pp. 2240–2248. doi: 10.1128/IAI.01585-07.

Osier, Faith Ha et al. (2014a) ‘Opsonic phagocytosis of Plasmodium falciparum merozoites: mechanism in human immunity and a correlate of protection against malaria.’, BMC medicine, 12(1), p. 108. doi: 10.1186/1741-7015-12-108.

Osier, Faith H. et al. (2014b) ‘Malaria: New antigens for a multicomponent blood-stage malaria vaccine’, Science Translational Medicine, 6(247). doi: 10.1126/scitranslmed.3008705.

Payne, R. O. et al. (2016) ‘Demonstration of the Blood-Stage Plasmodium falciparum Controlled Human Malaria Infection Model to Assess Efficacy of the P. falciparum Apical Membrane Antigen 1 Vaccine, FMP2.1/AS01.’, The Journal of infectious diseases, 213(11), pp. 1743–1751. doi: 10.1093/infdis/jiw039.

Payne, R. O. et al. (2017) ‘Human vaccination against RH5 induces neutralizing antimalarial antibodies that inhibit RH5 invasion complex interactions’, JCI Insight. doi: 10.1172/jci.insight.96381.

Persson, K. E. M. et al. (2006) ‘Development and optimization of high-throughput methods to measure Plasmodium falciparum-specific growth inhibitory antibodies.’, Journal of clinical microbiology, 44(5), pp. 1665–1673. doi: 10.1128/JCM.44.5.1665-1673.2006.

Proietti, C. et al. (2020) ‘Immune Signature Against Plasmodium falciparum Antigens Predicts Clinical Immunity in Distinct Malaria Endemic Communities.’, Molecular & cellular proteomics: MCP, 19(1), pp. 101–113. doi: 10.1074/mcp.RA118.001256.

Reiling, L. et al. (2019) ‘Targets of complement-fixing antibodies in protective immunity against malaria in children’, Nature communications. doi: 10.1038/s41467-019-08528-z.

Rono, J. et al. (2013) ‘Breadth of anti-merozoite antibody responses is associated with the genetic diversity of asymptomatic Plasmodium falciparum infections and protection against clinical malaria.’, Clinical infectious diseases: an official publication of the Infectious Diseases Society of America, 57(10), pp. 1409–1416. doi: 10.1093/cid/cit556.

Rohatgi, S. et al. (2013) ‘Fc Gamma Receptor 3A Polymorphism and Risk for HIV-Associated Cryptococcal Disease’, mBio. Edited by F. Dromer, 4(5), pp. e00573–13. doi: 10.1128/mBio.00573-13.

Stanisic, D. I. et al. (2009) ‘Immunoglobulin G subclass-specific responses against Plasmodium falciparum merozoite antigens are associated with control of parasitemia and protection from symptomatic illness.’, Infection and immunity, 77(3), pp. 1165–74. doi: 10.1128/IAI.01129-08.

Su, B. et al. (2019) ‘Update on Fc-Mediated Antibody Functions Against HIV-1 Beyond Neutralization.’, Frontiers in immunology, 10, p. 2968. doi: 10.3389/fimmu.2019.02968.

Teo, A. et al. (2016) ‘Functional Antibodies and Protection against Blood-stage Malaria.’, Trends in parasitology. England, 32(11), pp. 887–898. doi: 10.1016/j.pt.2016.07.003.

Tiendrebeogo, R. W. et al. (2015) ‘Antibody-dependent cellular inhibition is associated with reduced risk against Febrile malaria in a longitudinal cohort study involving Ghanaian children’, Open Forum Infectious Diseases. doi: 10.1093/ofid/ofv044.

Winkler, E. S. et al. (2021) ‘Human neutralizing antibodies against SARS-CoV-2 require intact Fc effector functions for optimal therapeutic protection.’, Cell, 184(7), pp. 1804–1820.e16. doi: 10.1016/j.cell.2021.02.026.

World Health Organization (2015) Guidelines for the treatment of malaria- 3rd edition. Geneva.

