## Supplementary for "Breadth of Fc-mediated effector function delineates grades of clinical immunity following human malaria challenge"

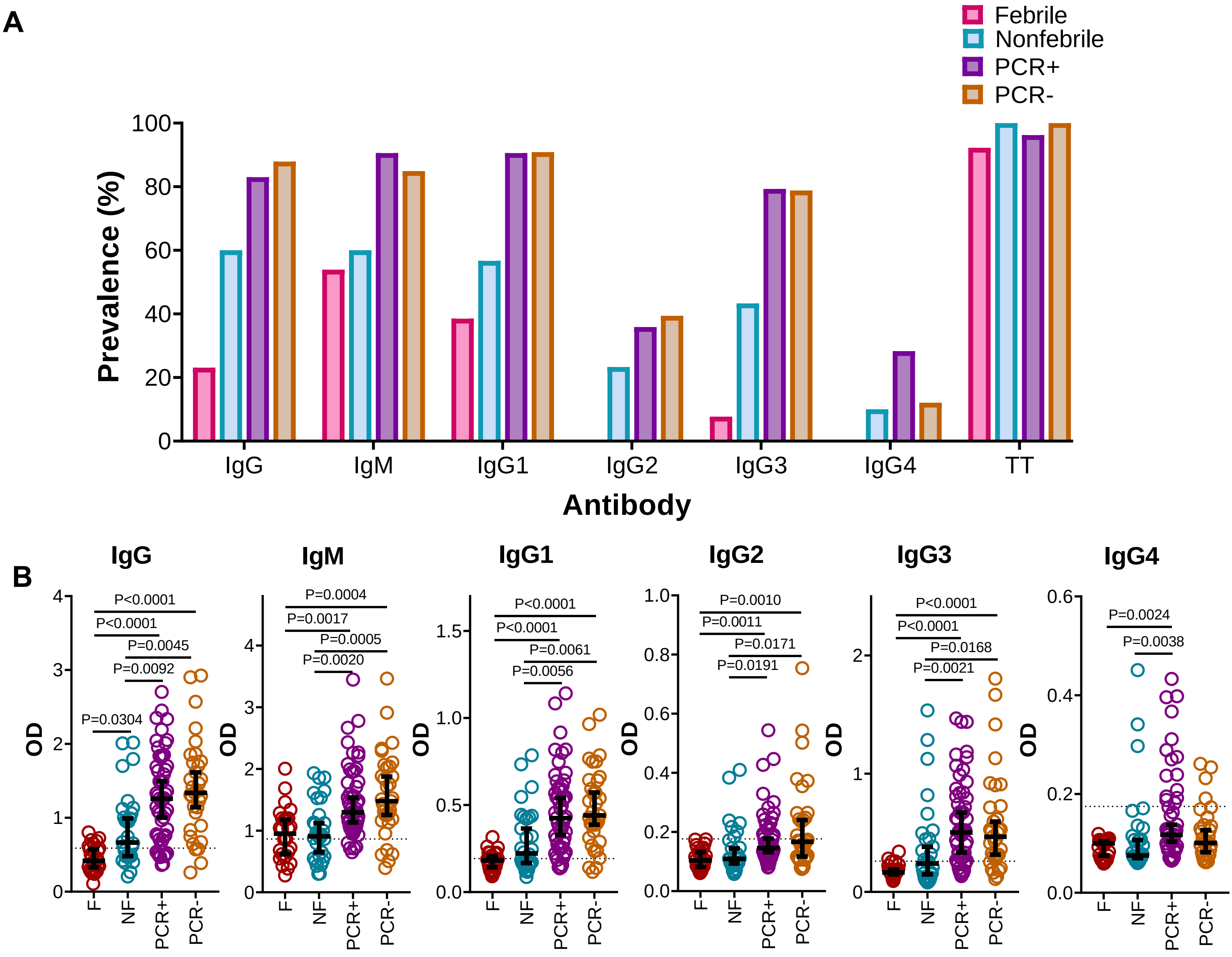


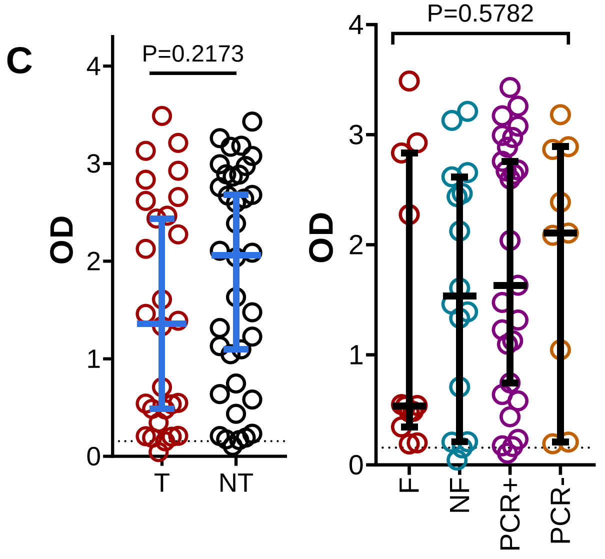


**Figure S1: Protection was associated with high levels of antibodies against merozoites**

(**A**) The prevalence of IgG, IgM, and IgG 1-4 antibodies against merozoites and total IgG against tetanus toxoid antibodies compared across the four phenotypes based on parasite growth patterns, treated febrile (F, n=26), treated non-febrile (NF, n=30), not treated PCR positive (PCR+, n=53) and not treated PCR negative (PCR-, n=33). (**B**) IgG, IgM, and IgG 1-4 antibody levels against merozoites compared across the four phenotypes based on parasite growth patterns. (**C**) Total IgG against tetanus toxoid were compared between treated (n=56) and not treated (n=86) and across the four phenotypes based on parasite growth patterns. Error bars represent median and 95% confidence intervals. P values were calculated using Mann-Whitney test for treatment outcome and using Kruskal Wallis test with Dunn's multiple comparisons test for the different phenotypes. The dotted black horizontal line represents the seropositivity cut-off (mean + 3SD of malaria naïve plasma samples).

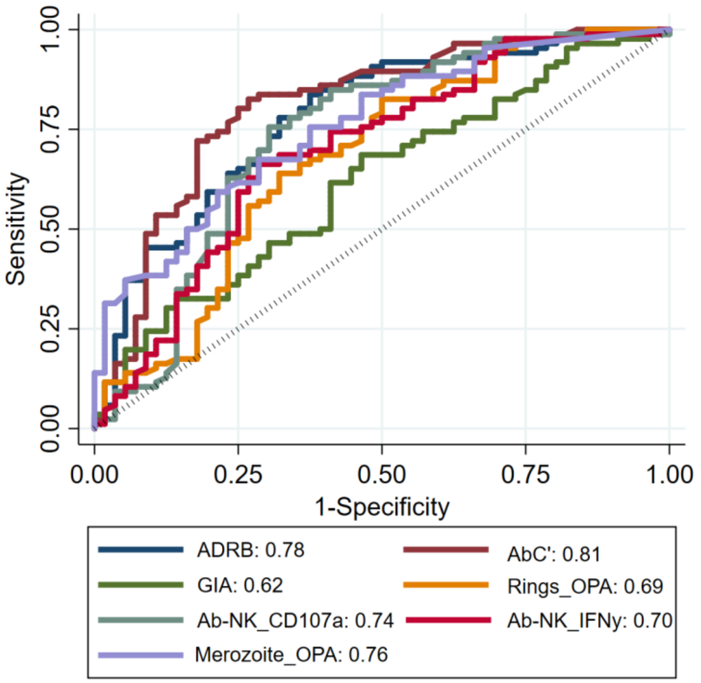


**Supplementary Figure S2: Fc-mediated effector functions were more strongly associated with protection than GIA**.

Receiver operating characteristic (ROC) curves for effector functions for all volunteers (n=142). The area under the ROC curve for each function is shown in the figure legend. ADRB; antibody dependent respiratory burst by neutrophils, AbC'; complement fixation activity, GIA; growth inhibition assay, Merozoite_OPA; opsonic phagocytosis of merozoites activity by monocytes, Rings_OPA; phagocytosis of ring stage parasites, Ab-NK_CD107a: antibody dependent granulation (CD107a) by natural killer cells, Ab-NK_IFNγ: antibody dependent IFNγ production by natural killer cells.

**Supplementary Table S1: Baseline characteristics of the CHMI-SIKA volunteers**

| | Cohort | | | Total |
|---|---|---|---|---|
| **Characteristic** | 2016 | 2017 | 2018 | |
| **Sample size** | 36 | 53 | 53 | 142 |
| **Age: median (range)** | 29 (18-44) | 25 (20-44) | 27 (18-45) | 27 (18-45) |
| **Sex: percentage male** | 75% | 73.6% | 60.4% | 69% |
| **No detectable lumefantrine** | 15 (41.6%) | 28 (52.8%) | 31 (58.5%) | 74 (52.1%) |
| **No detectable levels of any of the tested drugs[a]** | 13 (36.1%) | 23 (43.4%) | 28 (52.8%) | 64 (45.1%) |

[a] The following antimalarial drugs were measured in plasma samples collected on day 7 post challenge: Lumefantrine, pyrimethamine, sulfadoxine, chloroquine, artesunate and artemether. Of the 142 volunteers, 78 had detectable Lumefantrine or Sulfadoxine or both at levels below the minimum inhibitory concentrations (MIC, lumefantrine 200ng/ml, sulfadoxine 100ng/ml).

**Supplementary Table S2: Correlation of Fc-mediated effector functions with detectable lumefantrine and sulfadoxine drug levels**

| Fc-mediated effector functions | Lumefantrine (n=68) | | Sulfadoxine (n=10) | |
|---|---|---|---|---|
| | rho | *P* value | rho | *P* value |
| **GIA** | 0.58 | <0.0001 | 0.39 | 0.2632 |
| **ADRB** | 0.02 | 0.8739 | -0.04 | 0.9184 |
| **AbC'** | 0.19 | 0.1253 | -0.13 | 0.7330 |
| **Merozoites OPA** | 0.07 | 0.5779 | 0.00 | >0.999 |
| **Rings OPA** | 0.13 | 0.2992 | -0.14 | 0.7072 |
| **Ab-NK CD107a** | 0.11 | 0.3899 | -0.08 | 0.8382 |
| **Ab-NK IFNγ** | 0.07 | 0.5554 | -0.09 | 0.8113 |

ADRB; antibody dependent respiratory burst by neutrophils, AbC'; complement fixation activity, GIA; growth inhibition assay, Merozoite_OPA; opsonic phagocytosis of merozoites activity by monocytes, Rings_OPA; phagocytosis of ring stage parasites, Ab-NK_CD107a: antibody dependent granulation (CD107a) by natural killer cells, Ab-NK_IFNγ: antibody dependent IFNγ production by natural killer cells.

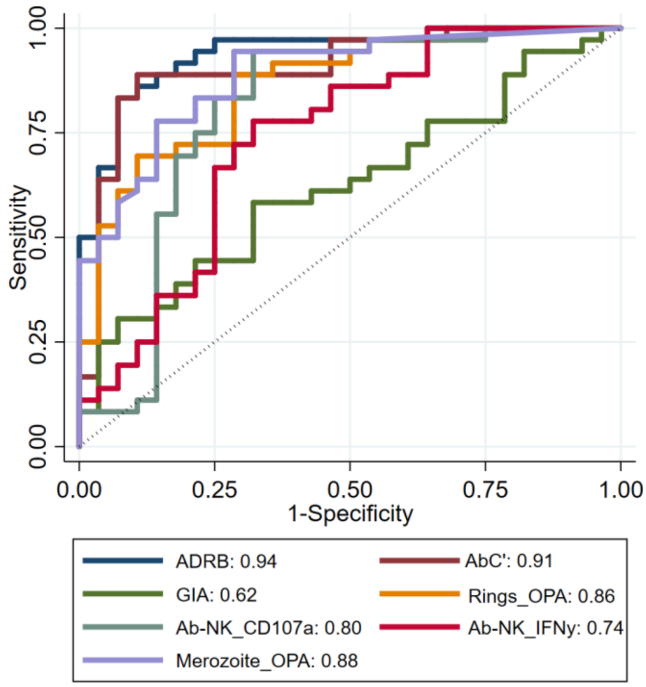


**Supplementary Figure S3: Fc-mediated effector functions were more strongly associated with protection than GIA**. Receiver operating characteristic (ROC) curves for effector functions for only drug-negative volunteers (n=64). The area under the ROC curve for each function is shown in the figure legend. ADRB; antibody dependent respiratory burst by neutrophils, AbC'; complement fixation activity, GIA; growth inhibition assay, Merozoite_OPA; opsonic phagocytosis of merozoites activity by monocytes, Rings_OPA; phagocytosis of ring stage parasites, Ab-NK_CD107a: antibody dependent granulation (CD107a) by natural killer cells, Ab-NK_IFNγ: antibody dependent IFNγ production by natural killer cells.

**Supplementary Table S3: Correlation of GIA with secondary outcome variables in drug negative samples**

| Secondary outcome variables | Spearman's rho (n=64) | *P* value | 95% confidence interval |
|---|---|---|---|
| **Mean parasite density** | -0.19 | 0.128 | -0.42 to 0.06 |
| **Maximum parasite density** | -0.27 | 0.031 | -0.49 to -0.02 |

The mean parasite density was calculated is a geometric mean of parasite density observed between days 8.5 and 22 after challenge, excluding timepoints after treatment; the maximum parasite density is the highest parasite density observed between days 8.5 and 22 after challenge, excluding timepoints after treatment. Data for drug-negative volunteers, n= 64.

**Supplementary Table S4: Correlation of anti-merozoite Fc mediated effector functions with secondary outcome variables in drug negative samples**

| **Secondary outcome variables** | **ADRB** | | **AbC'** | | **Merozoites OPA** | | **Rings OPA** | | **Ab-NK_CD107a** | | **Ab-NK_IFNγ** | |
|---|---|---|---|---|---|---|---|---|---|---|---|---|
| | **rho** | ***P*** | **rho** | ***P*** | **rho** | ***P*** | **rho** | ***P*** | **rho** | ***P*** | **rho** | ***P*** |
| **Mean parasitemia** | -0.69 | <0.0001 | -0.60 | <0.0001 | -0.57 | <0.0001 | -0.52 | <0.0001 | -0.41 | 0.0008 | -0.30 | 0.016 |
| **Maximum parasitemia** | -0.62 | <0.0001 | -0.59 | <0.0001 | -0.57 | <0.0001 | -0.50 | <0.0001 | -0.28 | 0.023 | -0.16 | 0.001 |

Spearman's correlation. The mean parasite density was calculated is a geometric mean of parasite density observed between days 8.5 and 22 after challenge, excluding timepoints after treatment; the maximum parasite density is the highest parasite density observed between days 8.5 and 22 after challenge, excluding timepoints after treatment. ADRB; antibody dependent respiratory burst by neutrophils, AbC'; complement fixation activity, OPA; opsonic phagocytosis activity by monocytes, Ab-NK_CD107a: antibody dependent granulation (CD107a) by natural killer cells, Ab-NK_IFNγ: antibody dependent IFNγ production by natural killer cells. Data for drug-negative volunteers, n= 64.

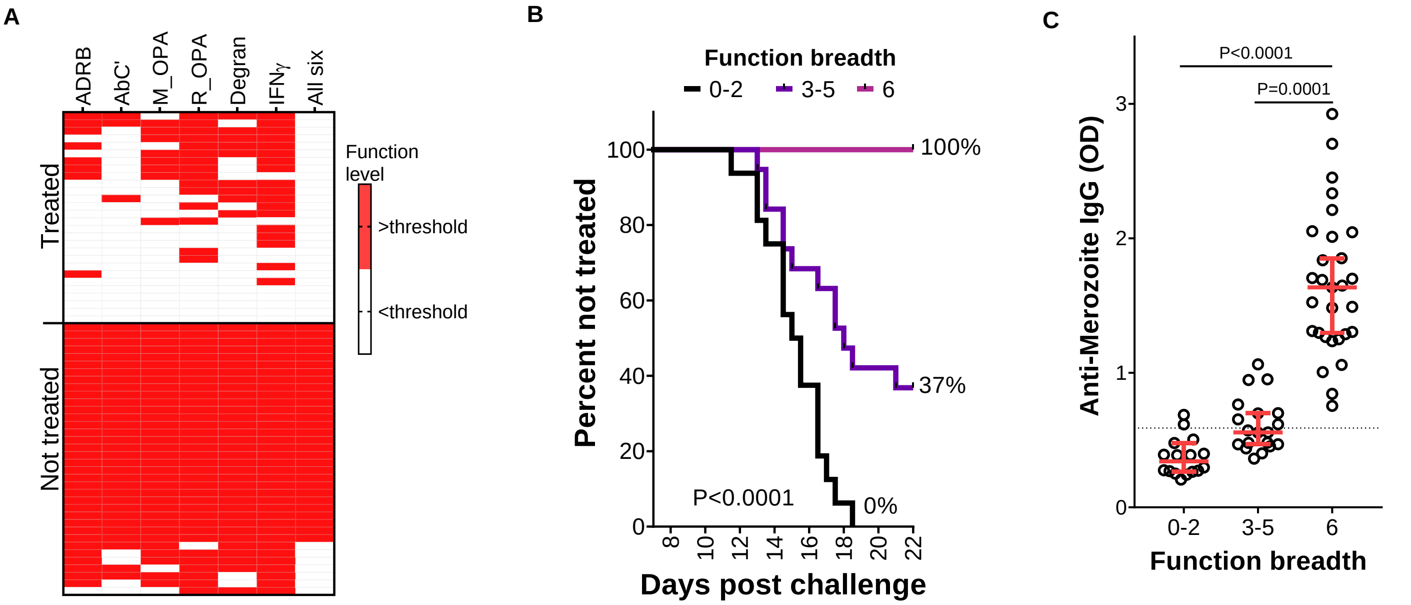


**Supplementary Figure S4: The breadth of Fc-mediated effector functions was more strongly associated with protection than any individual function.**

(**A**) A heatmap of all six Fc mediated effector functions in treated (n=28) and non treated (n=36) volunteers. Responses above function specific thresholds derived using maximally selected rank statistics are highlighted in red. Each column represents an Fc-mediated function while each row represents a volunteer. ADRB; antibody dependent respiratory burst by neutrophils, AbC'; complement fixation activity, M_OPA; opsonic phagocytosis of merozoites activity by monocytes, R_OPA; phagocytosis of ring stage parasites, Degran: antibody dependent granulation (CD107a) by natural killer cells, IFNγ: antibody dependent IFNγ production by natural killer cells. (**B**) Survival curves showing percentage of volunteers who remained untreated at different timepoints post challenge. Each line represents a function breath level starting with 0-2 (n=16), 3-5 (n=19) and 6 (n=29). The p value was calculated using Log-rank (Mantel-Cox) test. (**D**) Anti-merozoite IgG levels were compared between volunteers with varying breadth of Fc mediated function. Each dot represents an individual. Error bars represent median and 95% confidence intervals. P values were calculated using Kruskal Wallis test. The dotted black horizontal line represents the seropositivity cut-off (mean + 3SD of malaria naïve plasma samples). Data for drug-negative volunteers, n = 64.

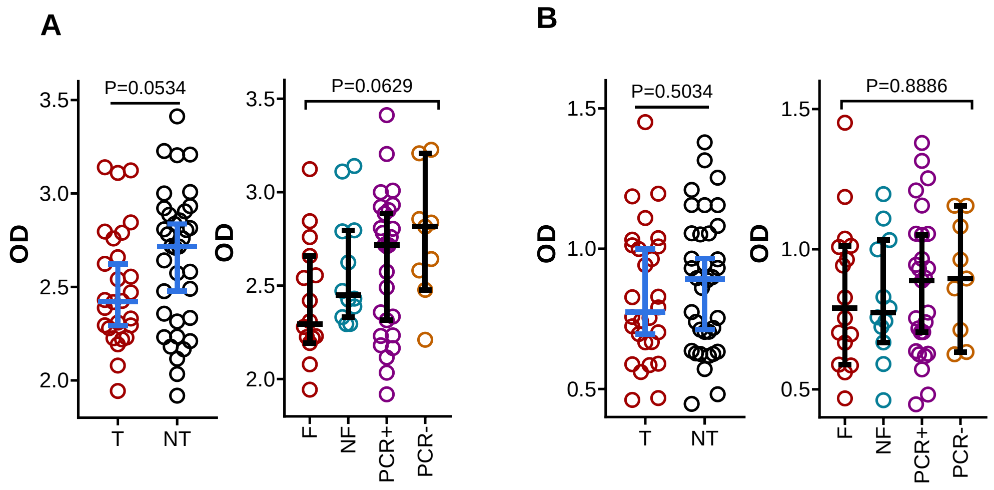


**Supplementary Figure 5: No significant differences in total antibody levels between protected and susceptible.**

(**A**)Total IgG and (**B**) total IgM antibodies were compared between volunteers who required treatment post challenge (n=28) and those who did not (n=36) and across the four phenotypes based on parasite growth patterns (Febrile n=16, non-febrile n=12, PCR positive n=27 and PCR negative n=9). Error bars represent median and 95% confidence intervals. p values were calculated using Mann-Whitney test for treatment outcome and using Kruskal Wallis test with Dunn's multiple comparisons test for the different phenotypes. Data for drug-negative volunteers, n = 64.
